# Targeting the Protein-Protein Interaction Between the CDC37 Co-Chaperone and Client Kinases by an Allosteric RAF Dimer Breaker

**DOI:** 10.1101/2025.02.17.637148

**Authors:** Alison Yu, Shrhea Banerjee, Sravani Malasani, Bamidele Towolawi, Zhiwei Liu, Zhihong Wang

## Abstract

Braftide, originally designed as a potent allosteric RAF kinase dimer disruptor, was intended to inhibit RAF dimerization by targeting the conserved RAF dimer interface. Intriguingly, Braftide has also been observed to trigger proteasome-mediated protein degradation with an unclear mechanism of action. This study elucidates the mechanism underlying Braftide’s dual functionality and assesses its potential as a chemical probe to target kinase-chaperone interaction. CDC37, a selectivity co-chaperone in the HSP90 chaperone machinery, plays a crucial role in facilitating the recognition of client kinase. The RAF dimer interface overlaps with the CDC37-kinase client recognition motif, known as the αC helix-β4 loop. Using co-immunoprecipitation and NanoBiT assays, we confirmed Braftide’s ability to selectively disrupt the CDC37-client kinase interaction while sparing HSP90. Through deuterium exchange mass spectrometry, molecular dynamic simulations, and *in vitro* crosslinking analyses, we mapped Braftide’s binding region within the BRAF kinase domain, as well as the CDC37 region implicated in the association of CDC37-client kinase complex. Consequently, this disruption destabilizes RAF kinase clients, resulting in proteasomal degradation, reduced cellular proliferation, and increased apoptosis in cancer cell lines. Furthermore, Braftide exhibits synergy with HSP90 inhibitors, jointly destabilizing both the CDC37-RAF complex and HSP90. Our work demonstrates the feasibility of disrupting the CDC37-client kinase interaction as an innovative therapeutic strategy and identifies the αC helix-β4 loop as a novel allosteric site with significant potential for the development of next-generation therapeutics.

## Introduction

The RAF kinase family, comprising ARAF, BRAF, and CRAF, is a core component of the MAPK (RAS-RAF-MEK-ERK) pathway, governing cell proliferation, survival, senescence, differentiation, and migration(1). BRAF, the most active RAF isoform, is dysregulated in 8% of all cancers and is intensely researched due to its role in oncogenesis(1). While ATP-competitive inhibitors like vemurafenib, dabrafenib, and encorafenib have shown success in treating BRAF V600E/K-mutant melanoma, they can lead to paradoxical activation of downstream ERK under certain conditions. This is linked to enhanced RAF dimerization, RAS-GTP association, and MEK binding(2–4), necessitating alternative therapeutic strategies.

To overcome these limitations, Braftide (TAT-miniPEG-TRHVNILLFM)(5), a peptide developed to disrupt RAF dimerization by targeting the conserved dimer interface among the RAF kinase family, along with other reported peptides (2, 6), demonstrate potent efficacy in downregulating the MAPK pathway. Braftide not only inhibits RAF kinase activity but also mitigates paradoxical activation induced by ATP-competitive inhibitors, both *in vitro* and in cells(5). The sequence of Braftide mirrors the residues spanning the tail-end of the αC helix and a region known as the αC helix-β4 loop, which includes three key residues: BRAF R509, L515, and M517, critical for RAF dimerization. Unexpectedly, Braftide induces protein degradation of RAF kinases, a novel mechanism reminiscent of PROTAC-mediated selective protein degradation. Targeting mutant BRAF using PROTACs has demonstrated degradation of mutant BRAF as a viable strategy to target the oncogenic kinase(7–9). The intriguing observation of protein degradation prompted us to investigate the underlying mechanism.

Protein degradation offers a promising approach to achieve sustained and robust inhibition of protein function. HSP90, an ATP-dependent protein-chaperone, is involved in the folding, maturation, and activation of client proteins, with selectivity directed by co-chaperones like CDC37(10). CDC37 is well established as a kinase co-chaperone that stabilizes kinase in a partially unfolded state(11). CDC37 is responsible for recognizing HSP90 clients by assessing the thermal stability of the client kinase(12). Subsequently, the CDC37-kinase domain complex engages with the HSP90 molecular chaperone(13). The CDC37-HSP90 complex ensures proper kinase folding and stability, including that of key oncogenic proteins such as BRAF, AKT, and HER2 (10, 12, 14). Disrupting this complex presents a novel therapeutic avenue. Inhibitors of HSP90 disrupt the HSP90 ATPase protein chaperone cycle, leading to proteasomal degradation of oncogenic proteins(14–16). However, direct targeting of HSP90 has proven challenging due to low selectivity and cytotoxicity, with over thirty ATP-competitive inhibitors failing to gain FDA approval (17, 18).

Recent structural studies of HSP90-CDC37-client kinase complexes suggest an alternative approach to selectively degrade oncogenic kinases by targeting the CDC37-client interaction, rather than directly inhibiting HSP90(16, 18, 19). Yet, to date, no inhibitors have been developed to efficiently disrupt this interaction. RAF family members are known clients of the HSP90-CDC37 chaperone complex and rely on their interaction with the CDC37-HSP90 chaperone complex to maintain basal protein stability(12). Prior biochemical studies, along with structures B/CRAF in complex with CDC37-HSP90, suggest that the αC helix–β4 loop (BRAF residues 510–520), a segment of the dimer interface (DIF), serves as the recognition site for CDC37(20–22). Although the HSP90 and CDC37 interface is vast and dynamic(15), the αC helix– β4 loop is a well-defined structural motif conserved across RAF and other kinases (23). The RAF DIF, being involved in CDC37-association, dimerization, and activation, emerges as an attractive therapeutic target. Our previous work demonstrated that Braftide, which inhibits the DIF, induces the degradation of BRAF/CRAF protein levels through the ubiquitin-26s proteasomal degradation system (UPS)(5). We propose that Braftide disrupts RAF-CDC37 interactions, thereby reducing RAF protein levels and offering a new strategy for targeting RAF-driven oncogenesis.

This study investigates Braftide’s ability to modulate the interaction between CDC37 and RAF kinases, exploring its role in promoting RAF degradation. We employ a combination of *in vitro* and cellular assays to demonstrate that Braftide destabilizes RAF through the ubiquitin-proteasome system and impairing cancer cell survival. Our findings suggest that disrupting the CDC37-client kinase interaction represents a viable alternative to direct HSP90 inhibition. Braftide’s ability to selectively disrupt the CDC37-RAF interaction, while sparing HSP90, provides an alternative and potentially more selective therapeutic strategy for targeting RAF and other oncogenic kinases.

## Results

### Braftide disrupts interaction between client RAF kinase and the CDC37 co-chaperone

A co-immunoprecipitation strategy was employed to monitor the association of BRAF and CRAF with CDC37 and HSP90 in the absence and presence of Braftide treatment. Our previous work revealed that Braftide treatment triggers proteosome-mediated degradation of BRAF and CRAF, therefore, all the cells were pre-treated with bortezomib before subjecting to Braftide (Figure S1A-G), unless noted otherwise (5). BRAF-FLAG or MBP-CRAF-FLAG (herein called CRAF-FLAG) was transiently transfected in HEK293 cells and immunoprecipitated for RAF’s association with the endogenous chaperone complex (Figure 1A-D). Notably, both Braftide-treated BRAF and CRAF reduced their association with CDC37 and HSP90 compared to the non-treated control (Figure 1A-D). Braftide effectively disrupts BRAF/CRAF’s association with the CDC37-HSP90 chaperone complex.

**Figure 1.**
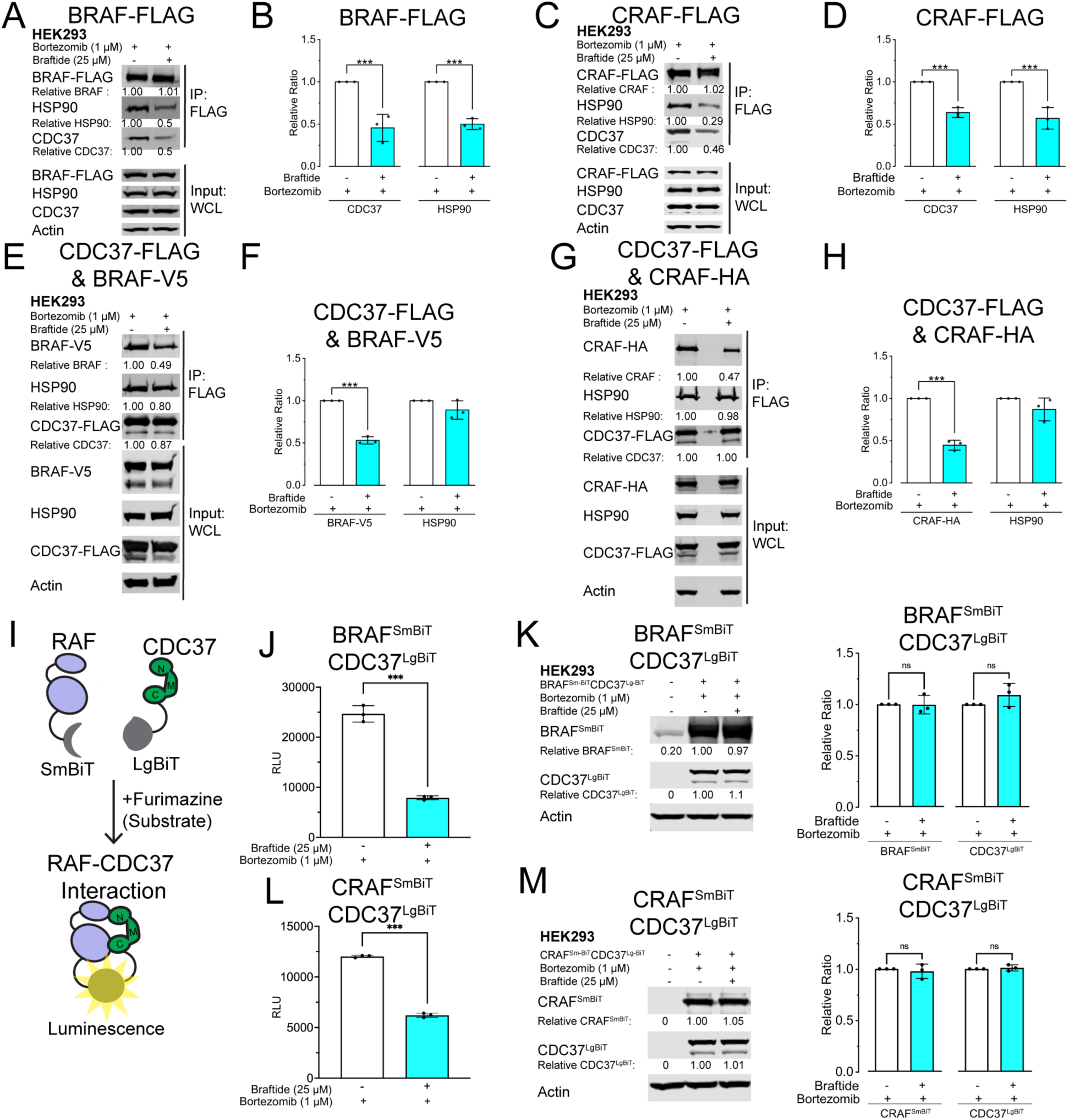
**Braftide disrupts interaction between client RAF kinase and the CDC37 co-chaperone**. A-D) BRAF-FLAG (A-B)/MBP-CRAF-FLAG (C-D) exogenously expressed in HEK293 cells in the absence and presence of 50 µM Braftide treatment (4 hrs). Representative immunoblot of immunoprecipitated (IP) B/CRAF-FLAG for the coimmunoprecipitated (co-IP) CDC37 and HSP90 chaperone complex. B, D) Densitometry analysis of CDC37 and HSP90 normalized to immunoprecipitated BRAF (B)/CRAF (D) across three biological replicates. E-H) CDC37-FLAG co-expressed with BRAF-V5 (E-F) or CRAF-HA (G-H). Representative blot of three biological replicates of immunoprecipitated CDC37-FLAG for co-IP BRAF-V5 (E) or CRAF-HA (G) and HSP90 in the absence and presence of Braftide treatment (25 µM) in HEK293 cells. I) Schematic illustrating NanoBiT luminescence assay. J, L) NanoBiT assay in the absence and presence of Braftide (25 µM, 4hrs) in HEK293 cells expressing NanoBiT constructs of BRAF^SmBiT^-CDC37^LgBiT^ (J) and CRAF^SmBiT^-CDC37^LgBiT^ (L). K, M) Representative immunoblot of HEK239 cells expressing BRAF^SmBiT^-CDC37^LgBiT^ (K) and CRAF^SmBiT^-CDC37^LgBiT^ (M) in the absence and presence of Braftide. K-M) Densitometry analysis of CDC37^LgBiT^ and BRAF^SmBiT^ (K) or CRAF^SmBIT^ (M) normalized to no treatment control across three biological replicates.

To further investigate whether this disruption specifically targets CDC37 or HSP90, we co-transfected CDC37-FLAG with BRAF-V5 or CRAF-HA in HEK293 cells and immunoprecipitated to assess their interaction with BRAF-V5 or CRAF-HA, and HSP90. Braftide treatment reduced the CDC37-client BRAF/CRAF complex while maintaining the CDC37-HSP90 interaction (Figure 1E-H). Similar results were observed in the KRAS-mutant colon cancer cell line HCT116, where Braftide reduced the CDC37-RAF interaction at lower doses than in HEK293 cells, suggesting this effect is not cell-line specific (Figure S1A-E).

Additionally, a NanoBiT assay was developed to monitor the CDC37-RAF interaction in live cells (Figure 1I). RAF and CDC37 are fused to two structurally complementary subunits of the NanoBiT enzyme, the small (Sm) and large (Lg) BiT. Structural complementation of the NanoBiT enzyme enables enzyme activity on furimazine, the substrate, and subsequently a luminescence-based readout of the CDC37 and RAF kinase interaction. In HEK293 cells co-expressing CDC37-LgBiT and RAF-SmBiT, luminescence indicated interaction between the two proteins. Braftide treatment led to a reduction in luminescence, reflecting a decrease in the CDC37-RAF interaction (Figure 1J, L), while proteins levels remained the same with bortezomib pre-treatment (Figure 1K, M). These results further confirm that Braftide disrupts the CDC37-RAF interaction in cells.

### Braftide directly binds to CDC37 and RAF, inducing proteasomal degradation of RAF via the ubiquitin-proteasome pathway

To identify proteins interacting with Braftide in cells, a crosslinking strategy was employed using biotinylated Braftide with a photo-active leucine analog, herein called BTN-Braftide, in HEK293 cells co-transfected with CDC37 and BRAF (Figure 2A). This photo-activatable leucine within BTN-Braftide contains a UV-responsive diazirine modification, enabling covalent bond formation with nearby amino acids upon UV exposure(24). Braftide contains three critical dimerization residues (BRAF aa R509, L515, M517) located in the RAF dimerization interface(5), conserved among all RAF family members (Figure 2B). The modified leucine in BTN-Braftide corresponds to residue 514 in the BRAF kinase domain, a position not involved in critical interactions in available structural data or in prior alanine scanning of the dimer interface (DIF)(6). Therefore, this alteration is not expected to affect the binding profile of Braftide. After UV light exposure, cells were collected, and the lysates were subjected to immunoprecipitation using streptavidin magnetic resin, followed by probing for BRAF, CRAF, CDC37, HSP90, and AKT1. BTN-Braftide co-immunoprecipitated CDC37, BRAF, and endogenous CRAF (Figure 2C). AKT1 is a well-established client kinase of CDC37(12), thus was selected as a control. Western Blots results demonstrate that neither AKT1 nor HSP90 was pulled down in this assay. Our results also demonstrate that BTN-JAK2, a control peptide derived from the αC helix-β4 loop of the JAK2 kinase, was used to treat HEK293 cells co-transfected with CDC37 and RAF. No protein degradation or reduction in pERK level was observed with this peptide (Figure S2A). Consequently, this peptide served as a scrambled peptide control and did not co-immunoprecipitate BRAF, CDC37, or HSP90 under the same conditions (Figure S2B). Together, these data suggest that Braftide selectively interacts with CDC37 and RAF.

**Figure 2.**
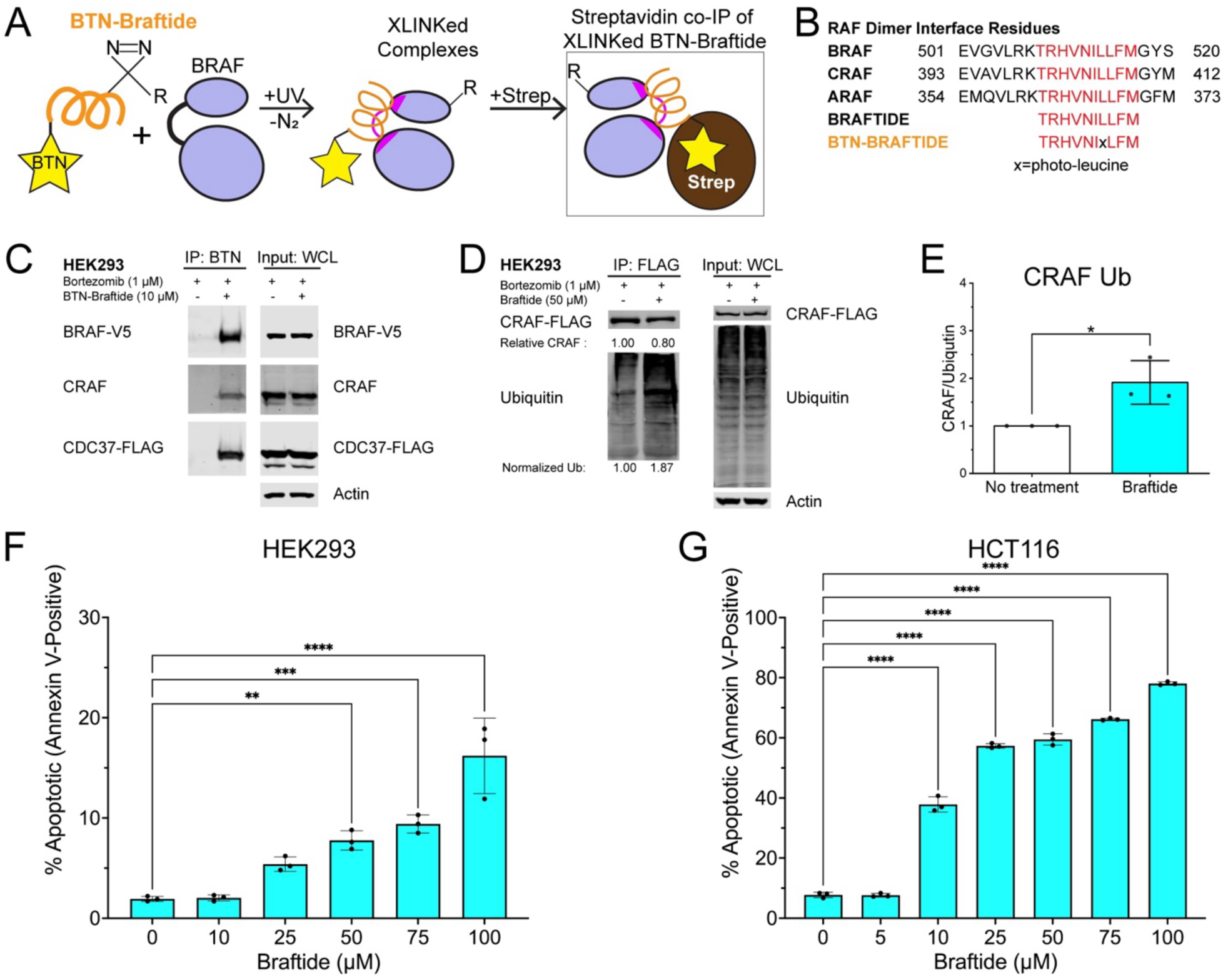
Braftide binds to CDC37 and RAF kinases and induces proteasomal degradation of the client kinase via the ubiquitin-proteasome pathway. A-B) Cartoon schematic of photo-activated crosslinking (XLINK) strategy of biotinylated (BTN)-Braftide (B) with BRAF. Crosslinked BTN-Braftide complexes are coimmunoprecipitated via streptavidin magnetic resin for further analysis. C) Representative immunoblot of overexpressed BRAF-V5 and CDC37-FLAG in HEK293 cells in the absence and presence of 10µM BTN-Braftide then immunoprecipitated and probed for individual proteins. D-E) Representative immunoblot (D) and densitometry (E) of overexpressed MBP-CRAF-FLAG immunoprecipitated and probed for ubiquitination in HEK293 cells in the absence and presence of 1 µM Braftide. F-G) Braftide mediated CDC37 disruption triggers apoptosis in HEK293 (F) cells and HCT116 (G) cells treated with indicated concentrations of Braftide for 4 hrs. Cells were stained with Annexin V, an apoptosis marker, and sorted via flow cytometry (n=3).

Dissociating the client kinase from CDC37 disrupts its interaction with the HSP90 chaperone. CRAF is reliant on its association with HSP90 and CDC37 for stability and function(12). Given that Braftide dissociates the CDC37-client interaction, we hypothesize that CRAF dissociation from CDC37 would increase CRAF instability, leading to its degradation via the ubiquitin-proteasomal pathway. This hypothesis was tested by monitoring CRAF ubiquitination in HEK293 cells expressing MBP-CRAF-FLAG, pre-treated with bortezomib to inhibit proteasomal degradation, both in the absence and presence of Braftide treatment. Braftide treatment led to increased CRAF ubiquitination (Figure 2D-E). These findings support our hypothesis that the dissociation of client kinase from the CDC37 co-chaperone destabilizes the client kinase, leading to subsequent degradation via the ubiquitin-proteasome pathway(5).

Previous studies have shown that knockdown of CDC37 expression via siRNA increases cellular apoptosis due to the degradation of client kinases in HCT116 and PC3 cells(25). Based on the observed increase in CRAF ubiquitination, we rationalized that Braftide-mediated dissociation of RAF kinases from CDC37 would similarly promote cellular apoptosis. HCT116 cells and HEK293 cells were subjected to increasing concentrations of Braftide (0, 5, 10, 25, 50, 75, and 100 µM) and stained for Annexin V, a marker of apoptosis. In both cell lines, Braftide triggered dose-dependent apoptosis by disrupting the CDC37-RAF interaction (Figure 2F-G, Figure S3). Notably, HCT116 cells showed higher sensitivity to Braftide compared with HEK293, likely due to their higher endogenous expression level of BRAF and CDC37 (Figure S4).

### Mapping the footprint of Braftide on BRAF kinase domain and CDC37

The crosslinking experiment (Figure 2C) captured the interaction between Braftide and BRAF-CDC37 in HEK293 cells. We used deuterium exchange mass spectrometry (HDX-MS) to identify specific interaction regions between Braftide and BRAF, providing insights into Braftide’s mechanism of disrupting RAF-CDC37 interaction. In these experiments, BRAF kinase domain purified from *E. coli* was incubated with and without Braftide, followed by exposure to deuterated buffer at specific time points (20s, 1min, 3min, 10min, 30min, 1.5hrs, 4.5hrs) to allow hydrogen exchange in solvent-accessible regions. After the exchange reaction, the samples were quenched, digested into peptides, and analyzed using mass spectrometry. Changes in deuterium incorporation between the Braftide-bound and unbound BRAF samples were quantified, with regions showing reduced deuterium uptake indicating potential binding sites covered by Braftide (Figure 3A). The entire BRAF kinase domain sequence was covered in this experiment (Figure S5), allowing comprehensive analysis of deuterium uptake differences, which were then mapped onto the BRAF structure (Figure 3B) to reveal specific interaction regions associated with Braftide binding. The identified binding region, spanning amino acids 496–515, overlaps with the BRAF dimer interface (Figure 3C), supporting the hypothesis that Braftide targets this interface in its role as a dimer breaker.

**Figure 3.**
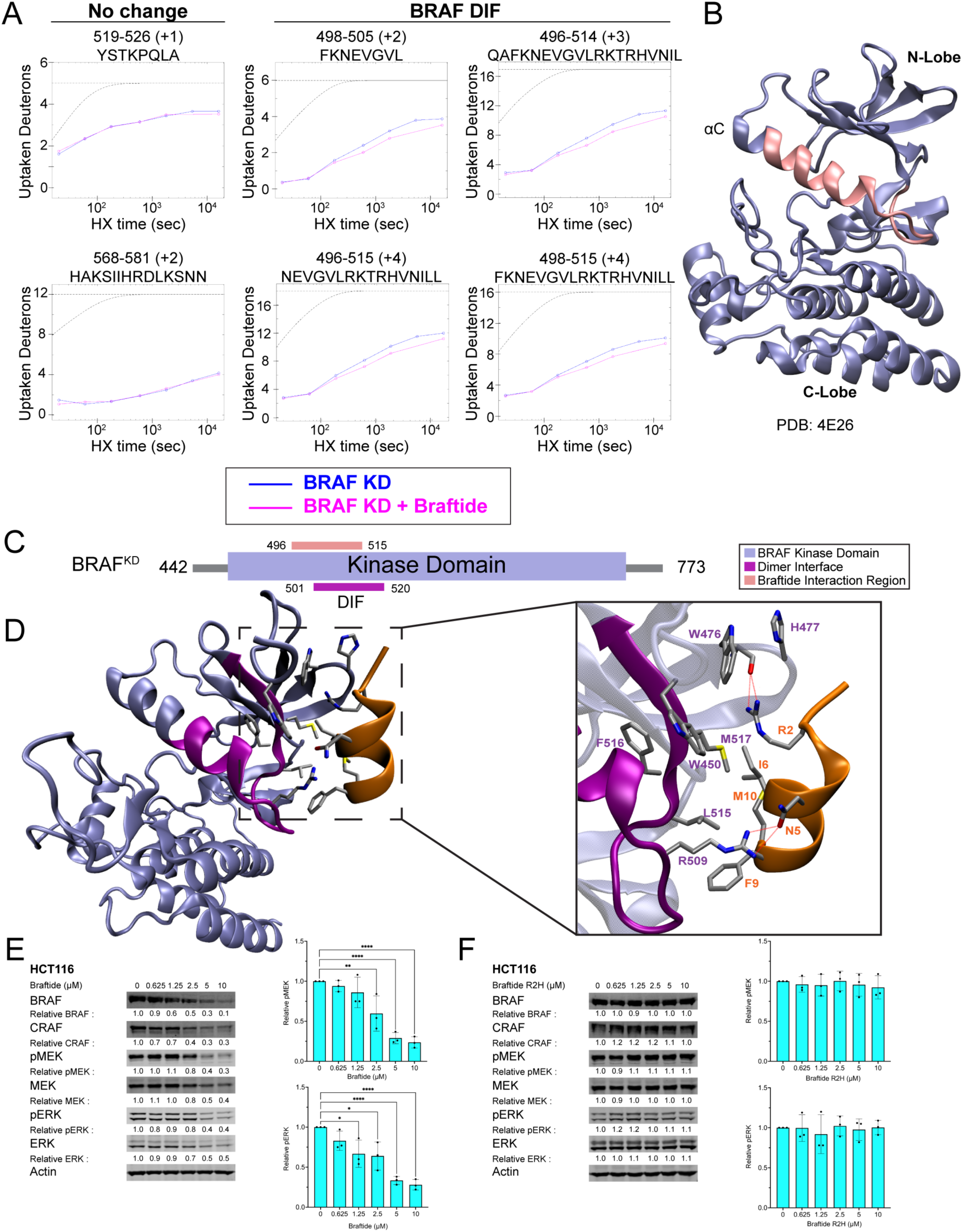
Braftide interacts with BRAF kinase domain via the αC helix-β4 loop of the dimer interface. A) Representative BRAF^KD^ peptides identified by hydrogen-deuterium exchange mass spectrometry (HDX-MS) in the absence (blue) and presence (pink) of Braftide. The peptide plots display slowed deuterium uptake, indicative of a broader trend across multiple overlapping peptides localized to the dimer interface. Gray dashed lines represent theoretical exchange profiles: the upper line corresponds to a fully unstructured peptide, while the lower line represents a peptide with complete hydrogen bond protection. B) Slowed deuterium uptake mapped onto the BRAF^KD^ crystal structure (PDB: 4E26), highlighting regions affected by Braftide (pink). C) Braftide interaction region on BRAF^KD^ (pink), according to HDX-MS, overlaps with the dimer interface (purple). D) Braftide (orange) binding to BRAF^KD^, MD simulations predicted interacting residues are shown in ball-and-stick. BRAF^KD^ crystal structure (PDB: 4E26), dimer interface region highlighted in purple. E-F) HCT116 cells were treated with Braftide (E) or Braftide^R2H^ mutant (F) at the indicated concentrations for 4 hrs. The cells were then harvested, lysed, and immunoblotted for MAPK proteins (BRAF, CRAF, MEK, phosphorylated MEK (pMEK), ERK, and phosphorylated ERK (pERK)). Densitometry analysis of pMEK and pERK normalized to (0-10 μM) Braftide (F) or Braftide^R2H^ (G).

In parallel, molecular dynamic (MD) simulations were applied to model the interaction between the BRAF kinase domain and Braftide, starting with the BRAF KD structure (PDB: 4E26) (26). The simulations indicated key interactions stabilizing the BRAF-Braftide complex, involving residues R509, L515, F516, and M517 from BRAF and R2, N5, I6, F9 and M10 from Braftide (Figure 3D, Figure S6A). These interactions include both hydrogen bonds and hydrophobic interactions. Those identified residues are part of the BRAF dimer interface, aligning with our HDX-MS data, which suggest that Braftide binds at this critical region.

Based on the structure of the CDK4 kinase in complex with the HSP90-CDC37, the αC helix-β4 loop has been suggested to serve as a potential molecular recognition motif for the CDC37 co-chaperone(26). By analyzing the structure of the BRAF-CDC37-HSP90 complex and overlaying the partially folded BRAF from this complex with a previously solved BRAF structure, we observed that the α helix-β loop of CDC37 superimposes on the αC helix–β4 loop of BRAF (Figure S6B). The α helix-β loop of CDC37 contains the same HxNI motif (residue numbers H20, P21, N22, I23) as the αC helix-β4 loop of RAF kinase (residue numbers H510, V511, N512, I513), suggesting that CDC37 applies a similar mechanism to engage with the C-lobe of protein kinase. This supports the idea that the αC helix–β4 loop in RAF is a potential site involved in mediating kinase-CDC37 interactions. This was further validated by the NanoBiT experiment, which showed that vemurafenib treatment, a competitive BRAF inhibitor known to stabilize the αC helix OUT configuration, has the potential to disrupt the interaction between BRAF and CDC37 (Figure S6C). MD simulations were conducted to identify the binding site of Braftide on CDC37 (Figure S7). These simulations revealed that Braftide binds to a region on CDC37 (residue numbers F29, M112, P113, W114) in close proximity to the previously identified client kinase recognition site (20-HPNI-23). To verify the interactions identified via MD simulations, we synthesized a modified R/H peptide, in which the R2 residue of Braftide, a key residue involved in interaction with CDC37 based on the simulation, was mutated to histidine. Unlike the unmodified Braftide (Figure 3E), the R/H peptide failed to induce protein degradation of BRAF, CRAF, MEK, and ERK, and had no impact on MAPK signaling in HCT116 cells (Figure 3F), verifying the key interactions observed in the MD simulations for maintaining the interaction between Braftide and CDC37. This result reinforces the proposed mechanism by which Braftide disrupts CDC37-mediated client stabilization.

### Braftide downregulates endogenous kinase levels via a structurally conserved αC helix-β4 loop interacting with CDC37

We propose that Braftide targets CDC37 by binding close to the region critical for client kinase recognition as shown in our MD simulations. Consequently, this interaction likely promotes the dissociation of CDC37 from a broader range of client kinases. This hypothesis is supported by the reduced association of CDC37 with AKT1, a well-characterized oncogenic kinase, upon Braftide treatment (Figure S8).

Our findings highlight the therapeutic potential of targeting CDC37-client kinase interactions, as oncogenic kinases are intrinsically unstable and thus are more dependent on their association with the HSP90-CDC37 chaperone complex for stability and activation compared with their wild-type counterparts (12). To assess the broader impact of Braftide on kinase levels, we selected HCT116 cells, which exhibit high endogenous expression of CDC37 and are more sensitive to Braftide treatment (Figure S4). HCT116 cells harbor an upstream KRAS G13C activating mutation, with a previously reported Braftide EC50 of 7.1 ± 0.5 µM(5). To investigate differential protein expression, three independent biological replicates of HCT116 cells were treated with or without 10 µM Braftide for 1 hr, followed by protein profiling using LC-MS/MS. Proteomic analysis identified 5,405 proteins (SI), of which 745 were differentially regulated with a cutoff of 2-fold change (FC) (Figure S8, S10). Among these, 84% (627 proteins) were downregulated (Figure S10A, neon green), encompassing proteins with functions such as catalytic activity, transferase, and hydrolase activity. Gene ontology (GO) analysis revealed that biological processes significantly impacted by Braftide included cellular and metabolic processes, signal transduction, and biosynthesis. While 16% (118 proteins) were upregulated, GO analysis indicated that catalytic activity and binding functions remained the most affected molecular functions among this subset (Figure S10B).

To validate the proteomic findings, several protein kinases identified as downregulated were selected for analysis via western blotting, including GSK3α, S6, AKT1, TAK, and FAK, along with MAPK pathway members BRAF, CRAF, MEK, and ERK. Consistent with the LC-MS/MS data (SI), treatment of HCT116 cells with 10 µM Braftide for 1 hr resulted in reduced protein expression levels of GSK3α, S6, AKT1, TAK, FAK, BRAF, CRAF, MEK1/2, and ERK1/2 (Figure 4A). In parallel, we examined the effects of Braftide treatment on several kinases identified as unchanged in protein levels by the proteomic analysis, including Chk2, PKCδ, and SRPK1. The results of these western blot experiments (Figure S11A) confirm that the protein levels of these kinases remain unchanged after Braftide treatment, demonstrating Braftide exhibits selectivity for CDC37-regulated kinases.

**Figure 4.**
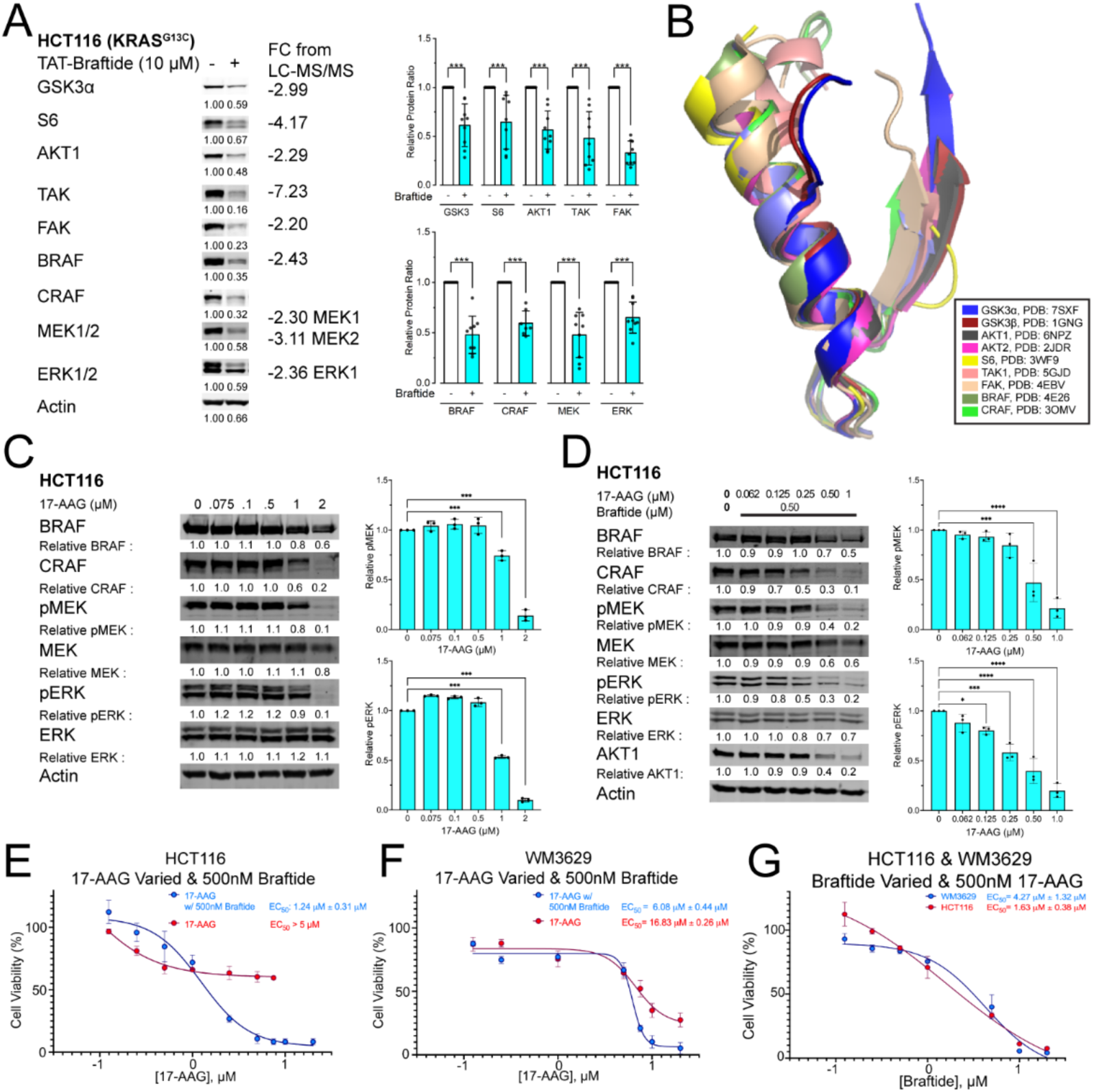
Braftide downregulates endogenous kinase levels and synergizes with HSP90 inhibitor 17-AAG. A) Representative immunoblot of selected kinases that are downregulated by Braftide treatment. Densitometry analysis of nine biological replicates of downregulated proteins (n=9). Graph bars represent the mean ± SD with corresponding P-values (*P<0.05, **P<0.01, ***P<0.001). B) Structural alignment of the conserved αC helix-β4 loop via Pymol, with indicated PDB accensions. C) HCT116 cells were treated with 17-AAG at the indicated concentrations for 16 hrs. The cells were then immunoblotted for MAPK proteins. Densitometry analysis of pMEK and pERK normalized to no treatment (0 μM) 17-AAG (n=3) D) HCT116 cells were treated with Braftide and 17-AAG at the indicated concentrations. Concentration of Braftide was held constant and 17-AAG concentrations were varied. The cells were then immunoblotted for MAPK proteins (n=3). Densitometry analysis of pMEK and pERK normalized to 0 μM 17-AAG and Braftide. E-G) Braftide and 17-AAG combination treatment inhibits cell proliferation in RAF dimer-dependent cancer cell lines, HCT116 (E) and WM3629 (F). E) HCT116 and F) WM3629 cells were treated with 17-AAG at the indicated concentrations (0, 0.125, 0.250, 0.500, 1.0, 5, 7.5, 10, and 20 μM) and a constant Braftide concentration (0.5 μM) for 24 hrs. Cells were treated with 17-AAG monotherapy at indicated concentrations for HCT116 (0, 0.125, 0.25, 0.5, 1, 2.5, 5, 7.5 μM) and same concentrations as combined therapy for WM3629. G) HCT116 and WM3629 cells were treated with Braftide at the indicated concentrations (0, 0.125, 0.25, 0.5, 1, 5, 10, 20 μM) and a constant 17-AAG concentration (0.5 μM) for 24 hrs. Cell viability was determined with the WST assay. EC50 values were obtained from a dose-response curve (4-parameter logistic equation).

The Braftide-mediated downregulation of kinase expression aligns with the decreased kinase levels (AKT, GSK3β, and CRAF) observed in siRNA mediated CDC37 knockdown in HCT116 cells(25). The αC helix-β4 loop, a conserved element typically 8 residues in length (with extensions in some kinases), anchors the N-and C-lobes through hydrophobic interactions and is one of the most stable structural elements in the kinase domain (28). The structural conservation within the αC helix-β4 loop region among protein kinases (Figure 4B) suggests that this motif may be a common interaction site between all client kinases and CDC37, serving as a recognition module for chaperone loading (Figure S6B). Taken together, these findings suggest that Braftide destabilizes the interactions between client kinases and the CDC37 co-chaperone, leading to a reduction in protein kinase levels in HCT116 cells. (26)

### Braftide synergizes with HSP90 inhibition to decrease cell viability

By targeting the structurally conserved αC helix-β4 loop, Braftide exhibits broad-spectrum effects on kinases that rely on CDC37 for stability and activation, providing a compelling basis for further exploration of CDC37 as a therapeutic target. Current efforts in HSP90 inhibition predominantly target ATP competitive inhibition to disrupt the chaperone cycle. However, this approach often requires high dosages, making cells more susceptible to thermal damage and inducing HSR1 expression(18). HSP90 inhibitors are limited by cytotoxicity arising from the inhibition of a broad and diverse client base. ATP competitive inhibitors for RAF kinase encounter key drawbacks such as paradoxical activation and drug resistance (29–31). Given CDC37’s role as a selectivity module for the HSP90 chaperone, we investigated whether disrupting the ternary complex via two approaches would synergize and promote cancer cell apoptosis. To overcome the limitations associated with RAF and HSP90 inhibitors, we propose inhibiting the ternary complex using Braftide in combination with sub-maximal concentrations of 17-AAG (17-N-allylamino-17-demethoxygeldanamycin), an HSP90 inhibitor. While 17-AAG is a potent small molecule HSP90 inhibitor, its clinical trials faced challenges due to high cytotoxicity(18). Leveraging Braftide’s capability to dissociate client kinase from the CDC37 co-chaperone, we speculate that 17-AAG could synergize with Braftide to reduce oncogenic kinase activity.

HCT116 cells were treated with 17-AAG alone or in combination with Braftide (Figure 4C-D). HSP90 monotherapy (17-AAG) depleted CRAF kinase level at a concentration of 2 µM (Figure 4C), while Braftide monotherapy(5) depleted RAF kinases levels at a concentration of 10 µM. To investigate potential synergy, we assessed combinatorial treatments in HCT116 cells, using a fixed concentration of 500 nM of the primary drug (Braftide) with varying concentrations (up to 1µM) of the secondary drug (17-AAG) (Figure 4D). Neither 17-AAG nor Braftide monotherapy exhibited obvious inhibition at the concentration of 500 nM. Intriguingly, the combination of 500 nM of 17-AAG and 500 nM Braftide resulted in significant degradation of CRAF kinase and downregulation of MAPK signaling, an effect unattainable with much higher concentrations of 17-AAG or Braftide alone. This clear synergy was evident at sub-maximal concentrations of both reagents, necessary to attenuate the MAPK pathway through disrupting the ternary complex (Figure 4D).

Based on the observed synergistic effects of inhibiting the ternary complex, the anti-proliferative effects were explored using combinatorial treatment in two RAF-dimer dependent cell lines: HCT116 and WM3629. HCT116 cells mainly rely on the wild-type BRAF-CRAF heterodimer and BRAF-BRAF homodimer to relay the MAPK signal, while WM3629 cells contain the BRAF D594G mutant, which is a kinase-dead mutant and depends on dimerization with CRAF for activation(27). The reported(27) EC50 of Braftide for WM3629 is 11.06 ± 1.12 µM. In HCT116 cells, the EC50 of 17-AAG in combinatorial treatment, with a constant Braftide concentration of 500 nM and varied 17-AAG concentrations (0, 0.125, 0.250, 0.500, 1.0, 5, 7.5, 10, and 20 μM), was determined to be 1.2 ± 0.31 µM (Figure 4E). In contrast, the EC50 value of 17-AAG in the absence of Braftide exceeded 5 μM (Figure 4E). Disrupting the CDC37-RAF complex in conjunction with 500 nM of Braftide resulted in an EC50 of 6.06 ± 0.44 µM for 17-AAG in mutant BRAF WM3629 cells, while the EC50 value of 17-AAG in the absence of Braftide approached 16.83 μM (Figure 4F). In both cell lines, we observed synergy between Braftide and 17-AAG. Additionally, in parallel, combination treatment with a constant 17-AAG concentration and varied Braftide concentration yielded EC50 values of 1.63 ± 0.38 µM and 4.27 ± 1.32 µM for HCT116 and WM3629 cells, respectively (Figure 4G). Annexin V staining was conducted to directly assess apoptosis in HCT116 cells treated with 17-AAG monotherapy and the combination of Braftide and 17-AAG. The results demonstrate a significant increase in apoptotic cell death with the combination therapy compared to monotherapy (Figure S11B-C). This effect is particularly evident at lower doses of the combination therapy, underscoring its enhanced apoptotic potential. Braftide exploits the reliance of oncogenic kinase dependence on CDC37 and synergizes with 17-AAG to sensitize cells to sub-maximal dosage treatment.

## Discussion

Our analyses have identified Braftide’s role in disrupting critical interactions within the cellular kinase regulatory network. Initially designed as an allosteric disruptor of RAF kinase dimerization (Figure 5A), our analyses uncover the multifaceted nature of Braftide, shedding light on its intricate mechanisms and therapeutic potential. Through HDX-MS, molecular dynamic simulations, proteomics analyses, structural elucidation, crosslinking, live cell NanoBiT assays, and co-immunoprecipitation, we uncover Braftide’s secondary function (Figure 5B). Braftide disrupts critical interactions for the stability and function of client kinases within the CDC37-HSP90 chaperone complex. By disrupting chaperone cycle entry, the client RAF kinase is destabilized, ubiquitinated, and degraded via the ubiquitin-proteasomal pathway. Our data collectively reveals the mechanisms underlying Braftide-triggered apoptosis in multiple cell lines, highlighting the therapeutic potential of targeting client kinase-CDC37 interactions in cancer cells.

**Figure 5.**
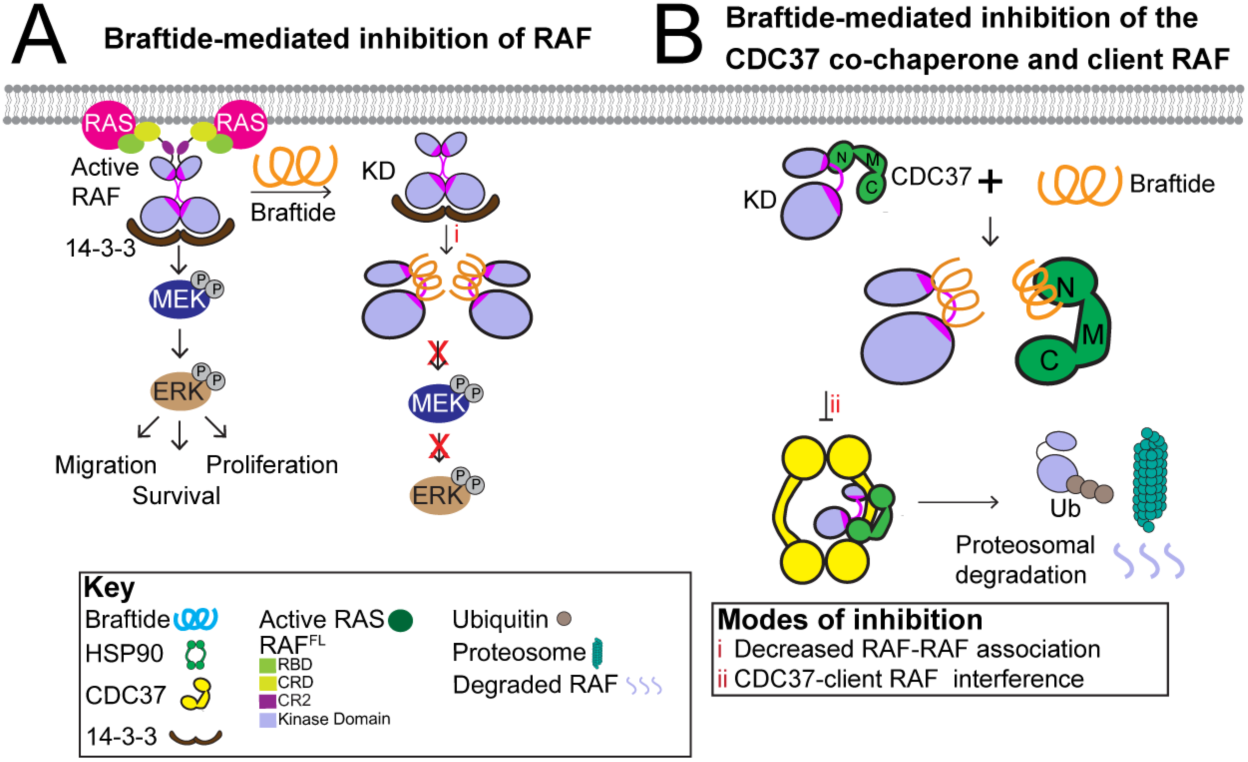
Braftide disrupts RAF dimers and the CDC37-RAF client kinase complex. A) Braftide binds to the dimer interface of RAF to dissociate RAF dimers. B) Braftide binds to CDC37 to disrupt the co-chaperone CDC37 with client kinases. Braftide treatment increases the ubiquitination of RAF, which leads to proteasomal-mediated degradation.

The αC helix-β4 loop is a conserved structural feature among eukaryotic protein kinases and has been suggested to coordinate the cooperativity between the N-lobe and C-lobe. In RAF kinases, the αC helix-β4 loop is a critical component of the dimer interface, essential for activation. In addition, the αC helix-β4 loop has been deemed as the recognition site of HSP90-CDC37 which is responsible for the proper folding of 60% of the human kinome. Structural analyses suggest that CDC37 mimics the conformation of the αC helix-β4 loop and uses the same mechanism to form hydrophobic interactions with the C-lobe of protein kinases, thereby leading to unfolding of the N-lobe of protein kinases and its subsequent loading into the lumen of HSP90 for folding. Hotspot mutations have been identified on the αC helix-β4 loop as well, highlighting the significance and versatility of this structural motif. Despite its central role in kinase function and regulation, the therapeutic potential of the αC helix-β4 loop has remained largely unexplored. Braftide is the first reported chemical tool that disrupts interactions between client kinases and the CDC37 co-chaperone. By mimicking the αC helix-β4 loop of RAF kinases, Braftide not only disrupts RAF dimerization but also impairs the interaction between kinases and CDC37. Our findings identify the αC helix-β4 loop as a key recognition site of CDC37, emphasizing its regulatory importance in kinase function folding and activation.

Our studies with Braftide demonstrate that therapeutic agents targeting the αC helix-β4 loop could selectively inhibit aberrant kinase signaling implicated in various diseases, including cancer and inflammatory disorders. Targeting this loop offers the potential for selective protein degradation of oncogenic kinases circumventing drug resistance mechanisms associated with traditional kinase inhibitors. By leveraging the principles of targeted protein degradation, braftide could serve as a prototype for designing molecules that promote the proteasomal clearance of dysregulated kinases driving oncogenesis. This also provides a great approach to target “undruggable” protein kinases such as pseudokinases, which rely on these interactions for function. As a pioneering tool, Braftide lays the foundation for further exploration of the αC helix-β4 loop as a therapeutic target. Future research into the structure, function, and evolutionary adaptations of this motif will provide deeper insights into kinase regulation and discovery of novel, highly specific protein kinase inhibitors.

Unlike conventional HSP90 inhibitors, Braftide preserves HSP90 activity while disrupting CDC37-client interactions, minimizing cytotoxicity and enhancing therapeutic precision. Braftide’s ability to destabilize RAF through CDC37 inhibition may offer an alternative to high-dose HSP90 inhibitors, mitigating their associated side effects. Conventional HSP90 monotherapies, such as 17-AAG, prove ineffective due to their impact on a large HSP90 client base, leading to global cellular dysregulation (16). While most HSP90-CDC37 inhibitors focus on HSP90 ATP-competitive inhibition(14), emerging strategies suggest targeting CDC37 to circumvent deleterious consequences of HSP90 inhibitors and indirectly inhibit the oncogenic kinase(16, 18). Current protein-protein interaction disruptors of the HSP90-CDC37 complex exclusively target HSP90 to allosterically dissociate the HSP90-CDC37 complex as observed by Conglobatin A and Platycodin D(34). While disrupting the HSP90 and CDC37 interface interferes with substrate client protein maturation, this mode of modulation also affects HSP90 activity(14, 35). Braftide emerges as the first chemical probe disrupting the CDC37-client kinase interface, preserving HSP90 activity, and synergizing with HSP90 inhibition to reduce cell proliferation. Our studies illustrate the promising effect of combining Braftide along with existing HSP90 inhibitors as a viable avenue to combat oncogenic kinase signaling by targeting the ternary complex and disrupting crucial kinase regulatory mechanisms. This dual inhibition minimizes promiscuity within large HSP90 client bases. These analyses form the foundation for refining peptides into small molecular inhibitor(s) targeting the CDC37 co-chaperone and dysregulated kinases for monotherapy and combined use with HSP90 inhibitors.

In conclusion, Braftide serves as both a powerful research probe and a prototype therapeutic agent, paving the way for targeted protein degradation strategies to combat kinase-driven diseases. Braftide represents a promising allosteric proof-of-concept disruptor of the CDC37-client kinase interaction and demonstrates the feasibility of targeting the association of client kinase with the chaperone complex. The development of peptide-guided small molecular inhibitors has demonstrated success, exemplified by a bespoke small molecule derived from the template macrocyclic peptide 1 to target nicotinamide N-methyltransferase for enzymatic inhibition in cells(36). Similar approaches should be employed to design CDC37-client small molecular disruptors, focusing on Braftide mimetics to enhance selectivity, stability, and cell permeability in these inhibitors. Future efforts aimed at transforming Braftide into a small molecule should prioritize enhancing selectivity for both the CDC37 co-chaperone and the oncogenic kinase. Mass spectrometry revealed a broad spectrum of kinases impacted by Braftide, suggesting Braftide interacts and inhibits CDC37 function beyond RAF kinase. Future studies should develop inhibitors that target specific pairs of CDC37-client kinase to achieve higher selectivity and reduced toxicity. Achieving a structure of Braftide in complex with BRAF and CDC37 will be instrumental in pursuing these objectives and guiding the development of more precise therapeutic intervention.

While this study emphasizes the impact of Braftide-targeted chaperone deprivation on RAF kinases, we acknowledge that similar effects are anticipated for other client protein kinases. This has been evidenced through our global proteomic analyses and cell-based assays. However, this does not diminish the significance of using Braftide as a chemical probe to explore the potential antagonism of oncogenic kinases beyond RAF kinases, aiming to achieve strong anti-tumor activities by depleting these oncogenic client protein kinases, such as CDK4, AKT, among others. Targeting CDC37 is believed to be superior to HSP90-directed inhibition, as it spares non-kinase HSP90 clients such as the steroid hormone receptors. While HSP90 plays diverse roles in various protein categories, CDC37 primarily functions in the biological functionality, stability, and regulation of protein kinases. The specificity underlying this preference is highly interesting yet remain largely unexplored due to the lack of suitable chemical probes.

## Materials and Methods

### Peptides

TAT-Braftide (GRKKRRQRRRPQ-miniPEG-TRHVNILLFM), BTN-Braftide (Biotin-GRKKRRQRRRPQ-miniPEG-TRHVNILxFM), TAT-Braftide^R2H^ (GRKKRRQRRRPQ-miniPEG-THHVNILLFM), TAT-JAK2 (GRKKRRQRRRPQ-miniPEG-SLQHDNIVKY), and BTN-JAK2 (Biotin-GRKKRRQRRRPQ-miniPEG-SxQHDNIVKY) where x is L-Photo-Leucine (ThermoFisher 22610)) was purchased from Lifetein with TFA removal. The purity was determined through HPLC (>95%) and confirmed through mass spectrometry.

### Plasmids

6X-HIS-BRAF-WT/FLAG or V5 was prepared as previously described(27, 37), and MBP-CRAF-FLAG was created using common cloning procedures with pcDNA^TM^ 4/TO (Invitrogen) as the vector. CDC37-myc-FLAG was purchased from Origene (RC201002).

### BRAF Kinase Domain (KD) Protein Purification

The BRAF kinase domain was purified as previously reported (38) with the human sequence expressed in *E. coli*.

### Hsp90:CDC37:BRAF KD Purification

Full-length mouse CDC37 and human BRAF KD were subcloned into the baculovirus vector pFastBac with a C-terminal hexahistidine tag on BRAF KD. *Sf9* cells were transfected with recombinant bacmid DNA for baculovirus production. Sf9 cells were infected for protein expression to obtain Hsp90:CDC37:BRAF KD construct. *Sf9* cell pellet was lysed by homogenization in lysis buffer (25mM Tris pH 8, 300mM NaCl, 10mM KCl, 10mM MgCl2, 2mM DTT, 10% Glycerol) with EDTA-free complete protease inhibitor tablets (Roche). Lysed cells were centrifuged to obtain supernatant of soluble cell lysate. Supernatant was incubated with pre-equilibrated cobalt resin for 2 h at 4°C. BRAF KD and CDC37 were co-eluted with endogenous *Sf9* HSP90 from TALON cobalt resin (Takara Bio) with increasing concentrations of imidazole. The complex was further purified on a Superdex 200 10/300 GL size exclusion chromatography column (Cytiva).

### Global Proteomic Identification of Braftide Regulated Proteins

HCT116 cells were treated in the absence and presence of 10 µM TAT-Braftide, 1 hr, 37°C. Cell pellets were harvested in PBS and cryopreserved. Samples were sent to be lysed and identified through mass spectrometry in three biological replicates (LC-MS/MS, The Wistar Institute, Philadelphia). Data was processed through the previously reported gene ontology analysis via the panther database(39) and plotted on Origin. For the volcano plot, the cutoff for the presented data is p=0.01 (**), q-value<0.05 (student’s t-test value adjusted for multiple comparisons using the Benjamini-Hochberg false discovery rate correction) and detected in at least 2 or more replicates. A total of 5405 proteins were identified, emphasizing 2-fold change in neon green and in red. Source data are available in the supplemental data files.

### Transient Transfection into mammalian cells

HEK293 cells were seeded at 1(10)^6^ cells per well for a 6 well plate of 5(10)^6^ for 10 cm dishes and incubated until 40-60% confluency. Transfection was carried out using a 1:3 DNA:PEI-MAX ratio in OPTI-MEM and allowed 48 hrs for protein expression. For peptide assays, 48 hrs post-transfection, cells were treated with TAT-Braftide carried in OPTI-MEM at indicated concentrations. Cells were harvested in cold PBS. Harvested cells were lysed in 4% SDS or modified RIPA for coimmunoprecipitation experiments. Quantification of total protein by the BCA assay was carried out according to the manufacturer’s protocols (Pierce 23225).

### Co-Immunoprecipitation in Cells

Transfected HEK293 cells were treated with indicated treatments and harvested in cold PBS. The cell pellets were lysed in modified RIPA buffer (50 mM HEPES pH 7.4, 150 mM sodium chloride, 0.1% NP40 (IGEPAL630), 1 mM EDTA, 5% glycerol, 1 mM phenylmethylsulfonyl fluoride, 20 mM beta-glycerophosphate, 2.5 mM sodium pyrophosphate, and 1 protease inhibitor tablet) and incubated with rotation for 2 hrs, 4°C. Cell lysates were then incubated with FLAG-M2 magnetic (SIGMA M8823) resin for 2 hrs to bind the FLAG-tagged BRAF/CRAF. The samples were washed 5 times in a modified RIPA buffer and quenched in loading dye and 95°C, 5 mins. Samples were analyzed via immunoblot. Treatment of cells was indicated in the figure legends.

### Cloning of NanoBIT Constructs

NanoBiT® CMV MCS BiBiT vector, which contains a BRAF N-terminal fused with LgBiT and CRAF N-terminal fused with SmBiT, were purchased from Promega. CDC37 LgBiT and CRAF SmBiT was generated using the standard Gibson Assembly (NEB HiFi Assembly) using NanoBiT® BiBiT as the vector and following primers: 5’ CCACCTCCTCCGAGAGAAACCACACTGACATCCTTCTCA 3’, 5’ TTTTGCAGCTAGCGATCGCCATGGTGGACTACAGCGTG 3’, 5’ GGCGATCGCTAGCTGCAAAAAG 3’, 5’ GTTTCTCTCGGAGGAGGTG 3’, and CDC37 LgBiT and BRAF SmBiT was generated using the standard Gibson Assembly using NanoBiT® BiBiT as the vector and following primers: 5’ CTATAGGGCTAGCGATCGCCATGGCGGCGCTGAGCGGTG 3’, 5’ CCACCTCCTCCGAGAGAAACGTGGACAGGAAACGCACC 3**’**, 5’ GTTTCTCTCGGAGGAGGTGG 3’, 5’ GGCGATCGCTAGCCCTATAG 3’.

### NanoBiT Luminescence Assay

HEK293 cells were transiently transfected with BRAF^SmBiT^-CDC37^LgBiT^ or CRAF^SmBiT^-CDC37^LgBiT^ at 2 µg plasmid per well and incubated at 37°C. Twenty-four hours post-transfection, cells were trypsinized, pelleted, resuspended in OptiMEM+4% FBS, and re-seeded onto 96-well plate at 5(10)^4^ cells per well then incubated at 37°C overnight. Cells were pretreated with 1 µM bortezomib for 5 hrs then Braftide for 4 hrs at 0 and 25 µM. After treatment 10 µM furimazine substrate was added to each well and luminescence readings were taken at 470-480 nm using CLARIOstar plate reader. Statistical significance was determined through paired t test where indicated, followed by Tukey’s honest significant different post-hoc test with corresponding P-values (*P<0.05, **P<0.01, ***P<0.001). All graph bars represent the mean ± SD with individual data points per biological replicate with corresponding p-values.

### Crosslinking BTN-Braftide and BTN-JAK2 to HEK293 overexpressed CDC37 and BRAF

HEK293 cells were co-transfected with CDC37 (Flag-tagged) and BRAF (V5-tagged). Cells were harvested in cold PBS and pellets were lysed in modified RIPA buffer with rotation for 2 hrs, 4°C. 1mg of cell lysate was incubated with 0 or 10µM BTN-Braftide/JAK2, with rotation for 4 hrs, at 4°C. Samples were then exposed to 365nM UV light for 1 hr over ice. After UV treatment, samples were incubated with Pierce Streptavidin magnetic beads (Cat#88816) for 17 hrs at 4°C, to bind BTN-Braftide. The samples were washed 5 times in high salt wash buffer (20 mM HEPES pH 7.4, 500 mM NaCl, 5% glycerol) and quenched with 25 mM Biotin in loading dye and 95°C, 5 mins. Samples were analyzed via immunoblot.

### Apoptosis Assays

HEK293 cells and HCT116 cells were treated with either control or Braftide at the indicated concentrations for 4 hrs. After treatment, both the floating and adherent cell populations were collected and adjusted with PBS (final concentration: 1 × 10^5^ cells/mL). The cells were then mixed with the Annexin V (Cytex FCCH100108) reagent in a 5:1 ratio (100 μL of cells mixed with 20 μL of annexin V reagent) and allowed to stain in the dark, at room temperature for 15 min. The cells were then analyzed on a Guava® Muse® Cell Analyzer (Millipore).

### Hydrogen Deuterium Exchange Mass Spectrometry (HDX-MS)

BRAF^KD^ (10 µM) in a buffer solution (20 mM HEPES pH 7.4, 150 mM NaCl, 5% glycerol in H2O), in the presence and absence of Braftide (30µM), was mixed with deuterated buffer (20 mM HEPES pH 7.4, 150 mM NaCl, 5% glycerol in D2O) at a 1:5 volume ratio (v:v) to initiate hydrogen-deuterium exchange. The exchange times varied from 20 seconds to 4.5 hrs. The reaction was quenched by mixing the sample 1:1 (v:v) with cold quench buffer (100 mM phosphate, pH 2.4, 0.5 M TCEP, 3 M guanidium chloride), lowering the pH to 2.4. The quenched solution was digested using an immobilized pepsin column, and the resulting peptides were trapped and desalted on a C8 column (Higgins Analytical TARGA C8, 5 µm, 5 x 1.0 mm). Peptides were desalted for 3 mins at 0°C before being eluted at 8 µL/min using a gradient of 10% to 40% acetonitrile over 15 mins. The eluted peptides were passed through an analytical column (Higgins Analytical TARGA C8, 5 µm, 50 x 0.3 mm) and introduced into a THERMO Q-Exactive mass spectrometer via electrospray ionization. Peptides were identified through MS/MS analysis of non-deuterated samples, with data analyzed using SEQUEST (Thermo Proteome Discoverer) against a database containing sequences of BRAFKD, Braftide, pepsin, potential contaminants, and decoy proteins. Deuterated samples were analyzed using ExMS2 software.

### Cell Viability

HCT116 and WM3629 cells were seeded onto poly-lysine coated, clear-bottomed, 96-well plates at 15(10)3 cells per well and allowed to incubate for 24 hrs. Cells were treated with Braftide and 17-AAG mixtures at indicated concentrations for 24 hrs (37°C). The WST-1 cell viability assay was carried according to the manufacturer’s protocols (Roche 11644807001) for 4 hrs and absorbance readings were taken at 450 nM in a CLARIOstar plate reader. For the dose-response curves of Braftide and 17-AAG, the data was fit to the 4-parameter logistic equation (dose-response fitting equation) in Graph Pad Prism:

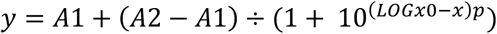

Error bars are indicative of the SEM of each point.

### Molecular Dynamics Simulations

Molecular dynamics simulation was performed using the Amber22 software package 1, with the FF19SB force field for protein 2. The initial structure of the BRAF KD-Braftide system was built based on the crystal structure of the active BRAF KD dimer (PDB ID: 4E26). Then the system was solvated in a periodic box with explicit OPC water molecules 3, and just the necessary number of counterions (Na + or Cl-) to neutralize the overall charge of the system. Equilibration of the system followed a consistent protocol which began with short minimization runs to fix possible bad contacts in initial structure, proceeded to gradually heating the system to 300K, then constant pressure (NPT) simulation at 1atm and 300K to equilibrate the simulation box size to be consistent with experimental density of an aqueous solution. Finally, a production run was performed in the NVT ensemble at 300K for 1μs, with a timestep of 1fs. The trajectories were then analyzed, focusing on the structural features and conformational changes at the BRAF KD-Braftide interface, with the cpptraj program in the Amber22 package.

### Immunoblot Analysis

Western blot band densities were quantified with ImageJ software (NIH) and normalized to the indicated loading control. Statistical significance was determined through one/two-way ANOVA where indicated, followed by Tukey’s honest significant different post-hoc test with corresponding P-values (*P<0.05, **P<0.002, ***P<0.0002, ****P<0.0001). All graph bars represent the mean ± SD with individual data points per biological replicate with corresponding p-values.

**Table 1.** Antibodies used in immunoblots.

| Manufacturer | Catalog Number | Name |
| --- | --- | --- |
| Sigma | F1804 | FLAG |
| Cell Signaling | 3724 | HA |
| Cell Signaling | 4694 | MEK1/2 |
| Cell Signaling | 9154 | pMEK |
| Cell Signaling | 4696 | ERK1/2 |
| Cell Signaling | 4370 | pERK |
| Cell Signaling | 2317 | S6 |
| Cell Signaling | 3700 | β-Actin |
| Cell Signaling | 4877 | HSP90 |
| Cell Signaling | 4222 | CDC37 |
| Cell Signaling | 3285 | FAK |
| Cell Signaling | 9422 | CRAF |
| Cell Signaling | 14814 | BRAF |
| Cell Signaling | 2938 | AKT1 |
| Cell Signaling | 4337 | GSK3α |
| Cell Signaling | 2289 | PP5 |
| Cell Signaling | 5206 | TAK1 |
| Cell Signaling | 3936 | Ubiquitin |
| Cell Signaling | 52012 | SRPK1 |
| Cell Signaling | 2662 | Chk2 |
| Cell Signaling | 2058 | PKCδ |
| Genetex | 42525 | V5 |
| Licor | 926-32211 | Anti-rabbit |
| Licor | 926-68070 | Anti-mouse |

### Inhibitors

Vemurafenib, 17-AAG (S1141) and bortezomib (S1013) were purchased from Selleckchem.

## Author contributions

AY, SB, and ZW were responsible for data analysis and manuscript preparation. AY, SB, SM, BT, and ZL were responsible for data generation. AY and ZW were responsible for original conceptualization.

## Supporting information

Mass spectrometry data from Braftide treated HCT116 cells with differential protein expression

## Acknowledgements

The authors acknowledge Dr. Susan Taylor (University of California, San Diego) for the insightful discussion.

The authors acknowledge Thomas Beer and Hsin-Yao Tang at the Wistar Institute’s Proteomics & Metabolomics Facility for performing the LC-MS/MS experiments and analysis and helpful discussions.

The authors acknowledge Robert Chitren and Subash Jonnalagadda at the College of Science and Mathematics, Rowan University for assistance with flow cytometric experiments and helpful discussions.

## Data Sharing Plan

All data is included in the manuscript and supporting information.

## Supplemental Information and Figures

**Figure S1.**
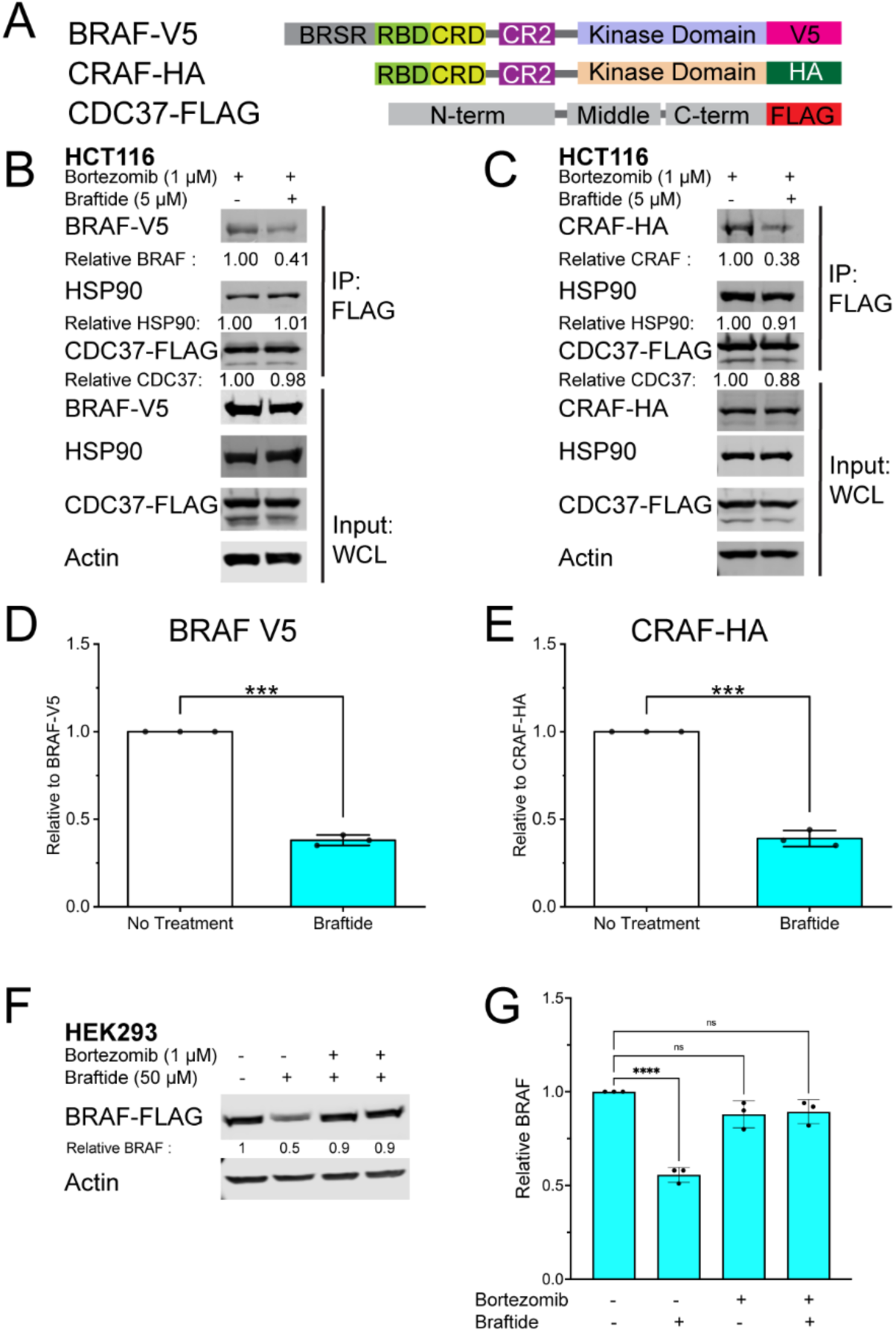
Braftide disrupts the CDC37 chaperone complex in HCT116 cells. A) Cartoon schematic of constructs used for probing the client RAF and CDC37 interaction B-C) Representative immunoblots of HCT116 cells exogenously expressing BRAF-V5 (B) or CRAF-HA (C) with CDC37-FLAG in the presence and absence of 5 µM Braftide. Immunoprecipitated CDC37-FLAG decreases its association with coimmunoprecipitated BRAF-V5 and CRAF-HA. D-E) Densitometry analysis of BRAF (D) and CRAF (E) association with CDC37 across three biological replicates. F) Representative immunoblot of overexpressed BRAF-FLAG in HEK293 cells treated with Braftide in the absence and presence of bortezomib. G) Densitometry analysis of BRAF levels in WCL across three biological replicates.

**Figure S2.**
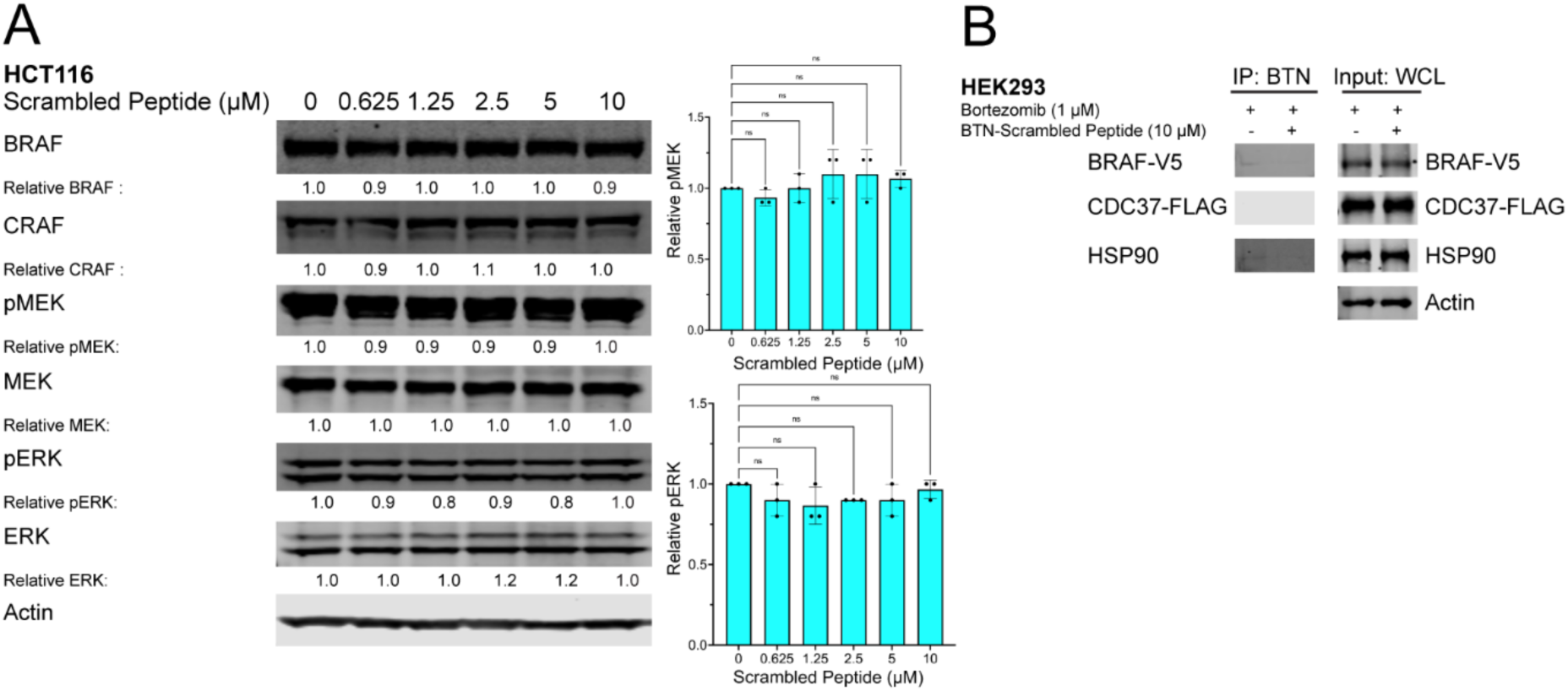
MAPK proteins are unaffected by JAK2 control peptide. A) HCT116 cells were treated with JAK2 control peptide at the indicated concentrations for 4 hrs. The cells were then harvested, lysed, and immunoblotted for MAPK proteins. Densitometry analysis of pMEK and pERK normalized to no treatment (0 μM) JAK2. B) Representative immunoblot of overexpressed BRAF-V5 and CDC37-FLAG in HEK293 cells in the absence and presence of crosslinked 10µM BTN-JAK2 then immunoprecipitated and probed for MAPK proteins.

**Figure S3.**
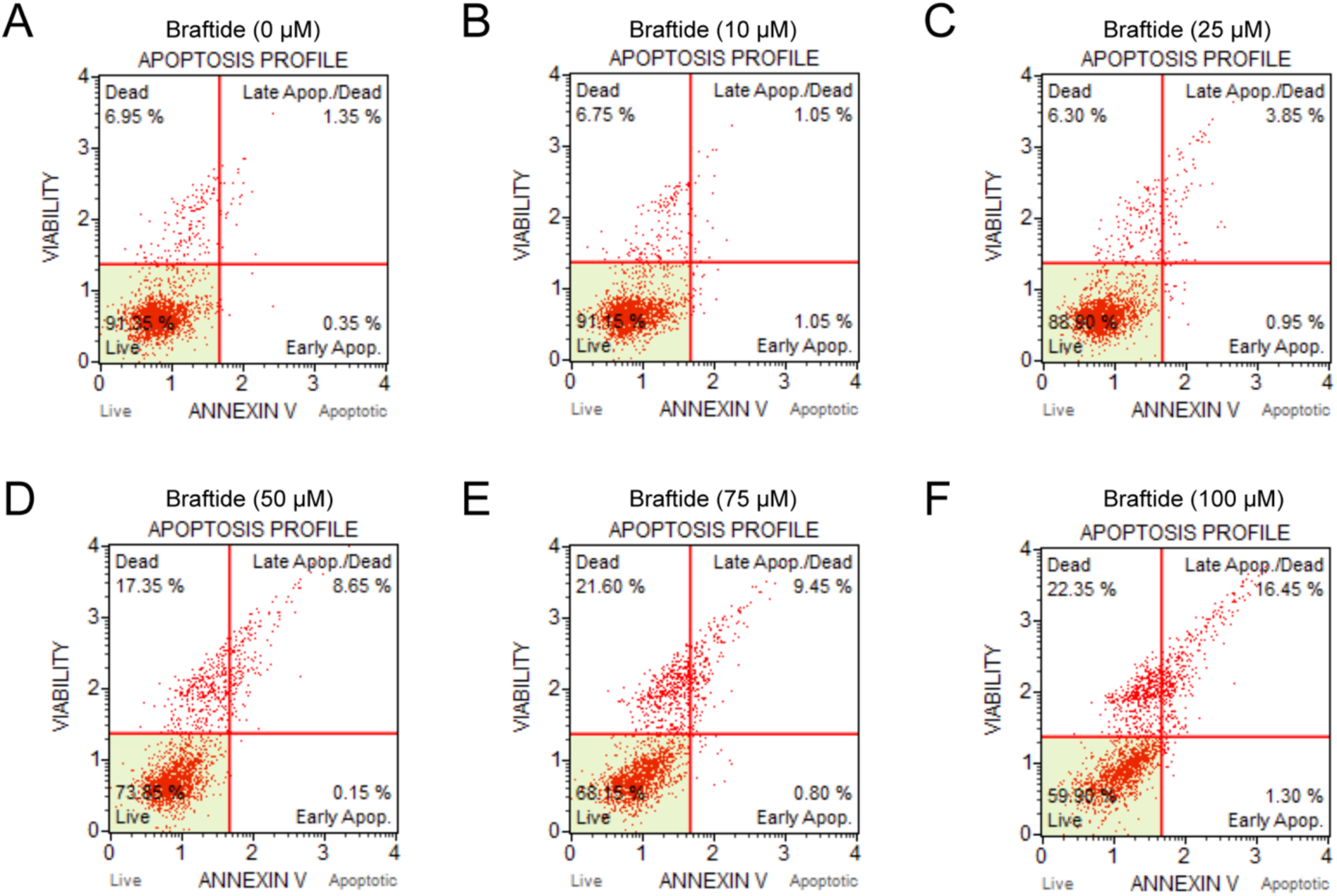
Braftide treatment triggers apoptosis. A-F) Representative apoptosis profile of HEK293 cells exogenously expressing WTFL BRAF treated with Braftide at 0 (A), 10 (B), 25 (C), 50 (D), 75 (E), and 100 µM (F) for 4 hrs.

**Figure S4.**
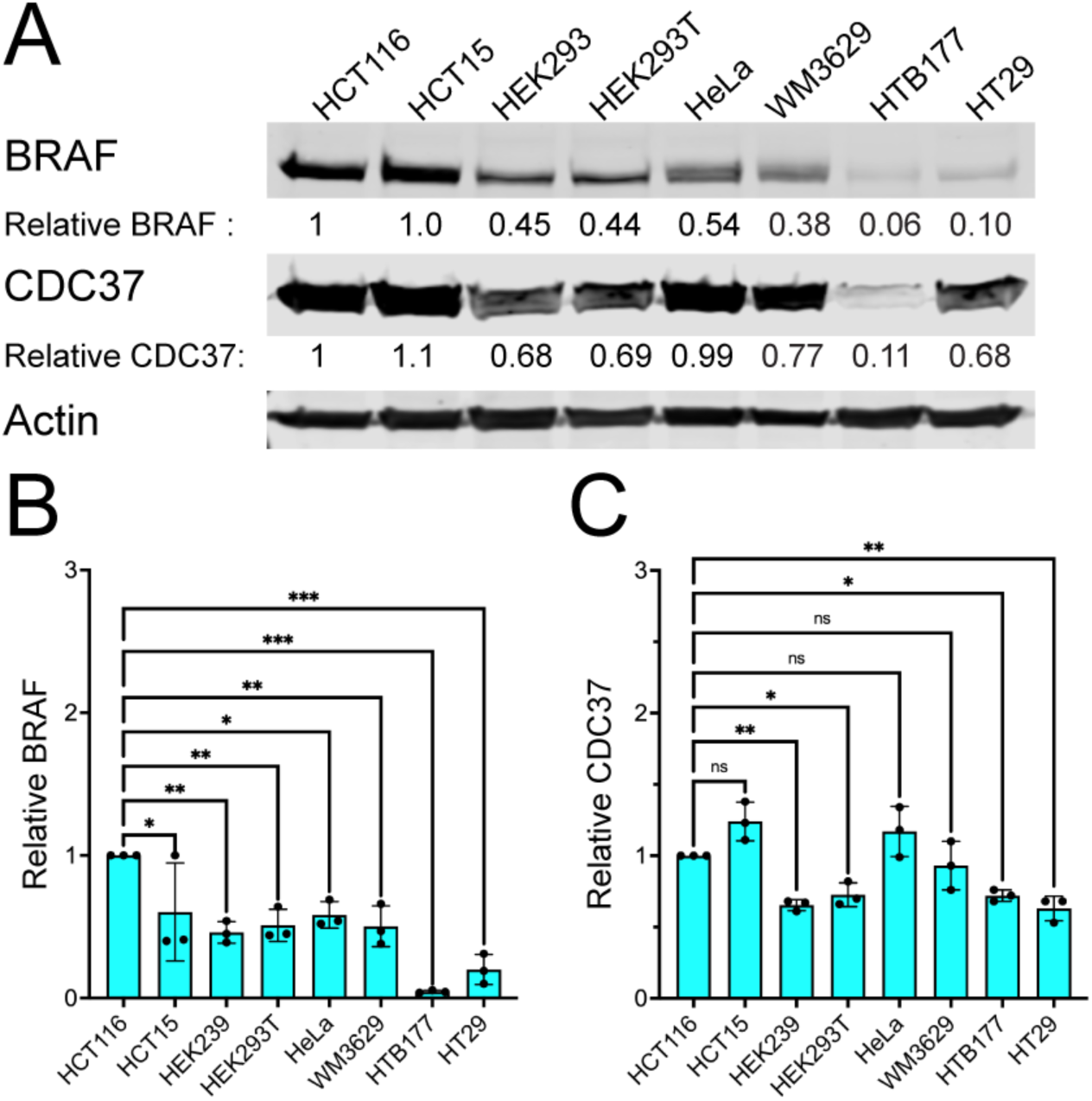
HCT116 expresses high BRAF and CDC37 across several cell lines. A) Representative immunoblot of several cell lines probed for endogenous BRAF and CDC37 across three biological replicates B-C) Densitometry analysis of BRAF (B) and CDC37 (D) across three biological replicates (n=3). Graph bars represent the mean ± SD with corresponding P-values (*P<0.05, **P<0.01, ***P<0.001). Statistical significance was determined via one-way ANOVA, followed by the post-hoc Tukey’s test.

**Figure S5.**
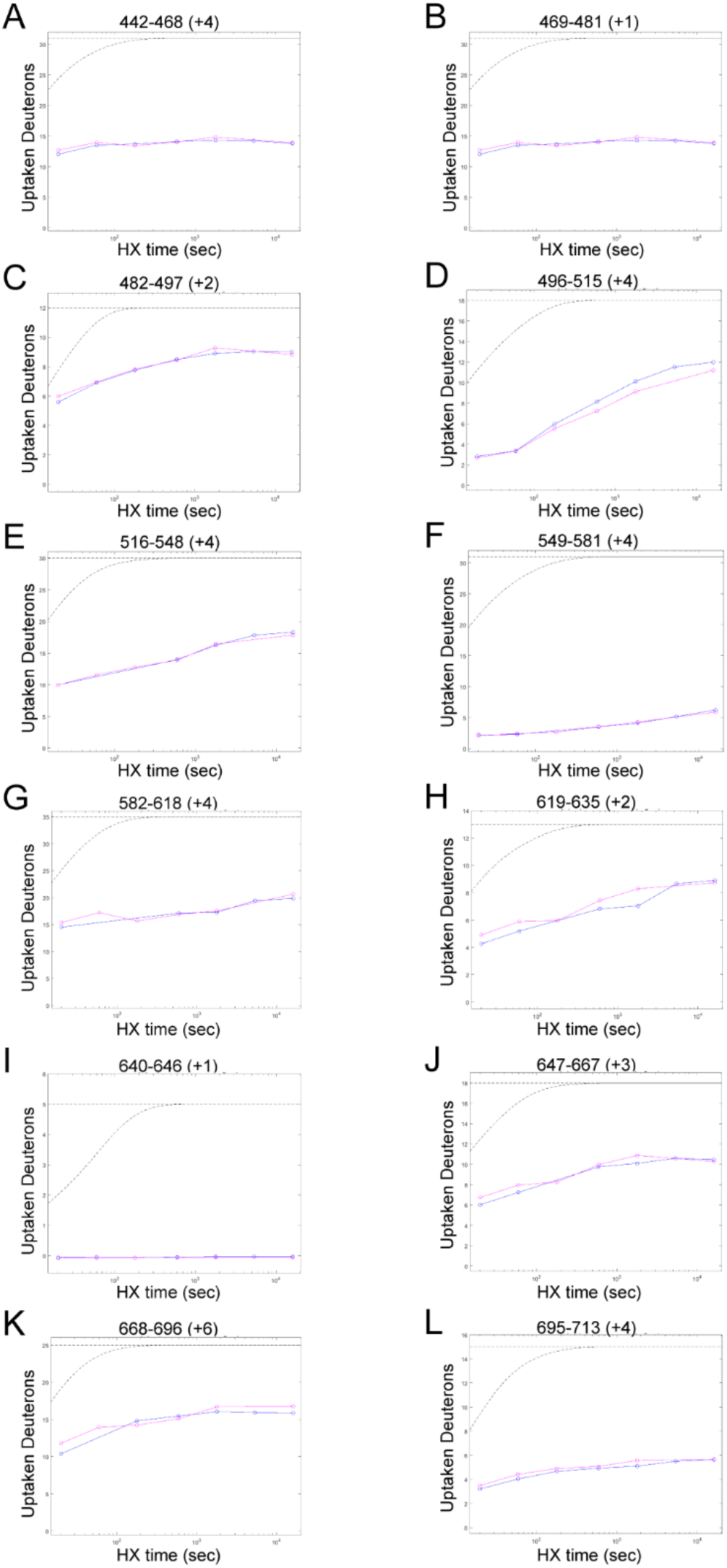
Braftide slows deuterium uptake in the DIF region of BRAF. A-L) Peptide plots spanning the entire BRAF^KD^ region that show essentially no change except in the dimer interface region, in the absence (blue) and presence (pink) of Braftide. Gray dashed lines represent theoretical exchange profiles: the upper line corresponds to a fully unstructured peptide, while the lower line represents a peptide with complete hydrogen bond protection.

**Figure S6.**
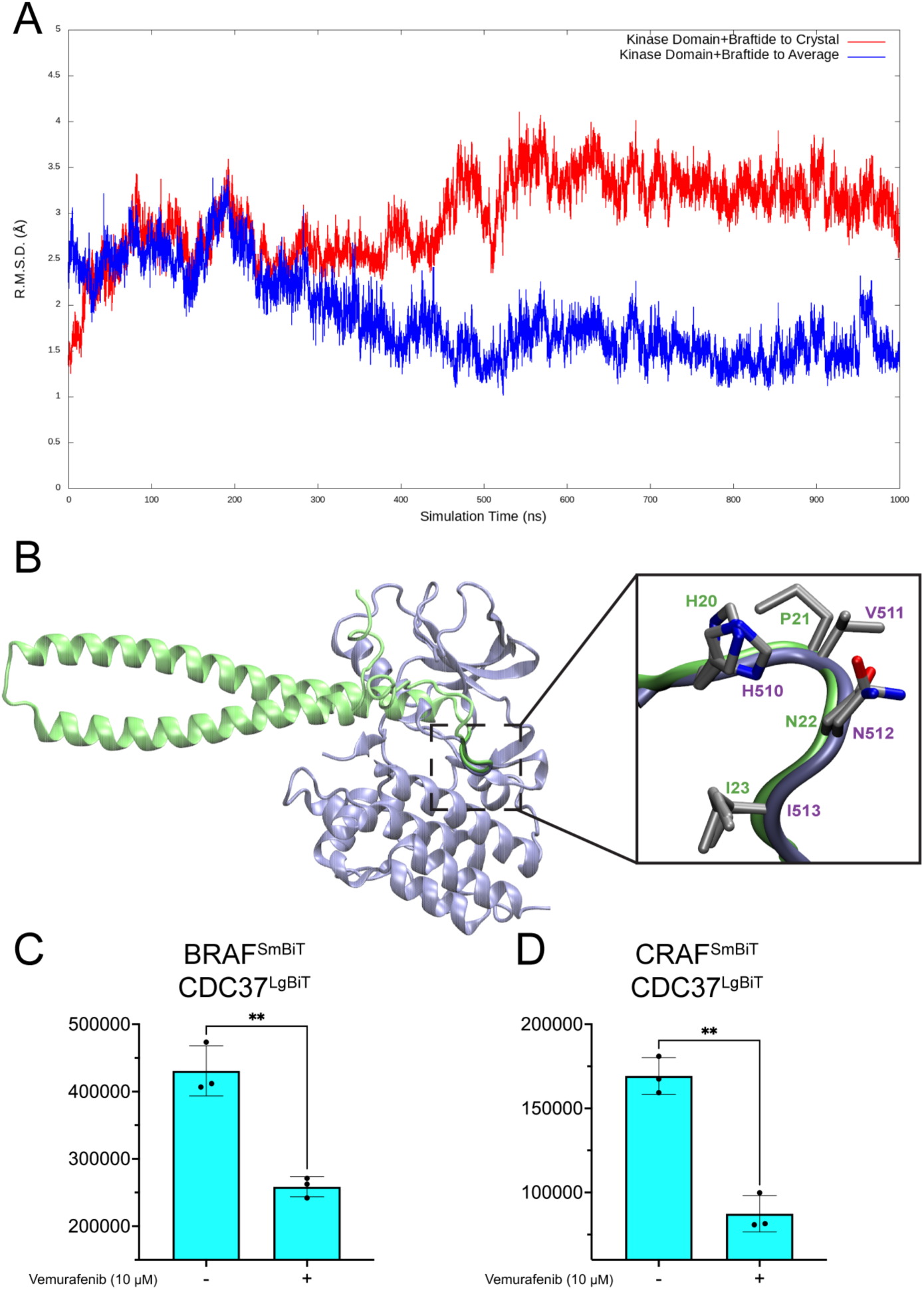
CDC37 shares the HxNI motif with client kinases. The binding dynamics between the BRAF^KD^ and the Braftide peptide, was performed via atomistic MD simulation in aqueous solution. A) Root Mean Square Displacement (RMSD) of Ca along the MD trajectory. The RMSD with respect to the crystal structure (red) starts at a low value of 1.5 Å and gradually increases and stabilizes roughly at around 3.5 Å from 500 ns, indicating conformational change from the crystal structure and possible convergence of structure. An average structure from 500ns to 1μs is obtained and RMSD with respect to this average structure are calculated (blue). This RMSD indicates a clear convergence of structure starting from 500 ns. B) Overlay of α helix-β loop of CDC37 (green) and αC helix-β4 loop of BRAF^KD^ (gray) with the HxNI overlapped C-D) NanoBiT assay in the absence and presence of Vemurafenib (10 µM, 4hrs) in HEK293 cells expressing NanoBiT constructs of BRAF^SmBiT^-CDC37^LgBiT^ (C) and CRAF^SmBiT^-CDC37^LgBiT^ (D) normalized to no treatment control across three biological replicates.

**Figure S7.**
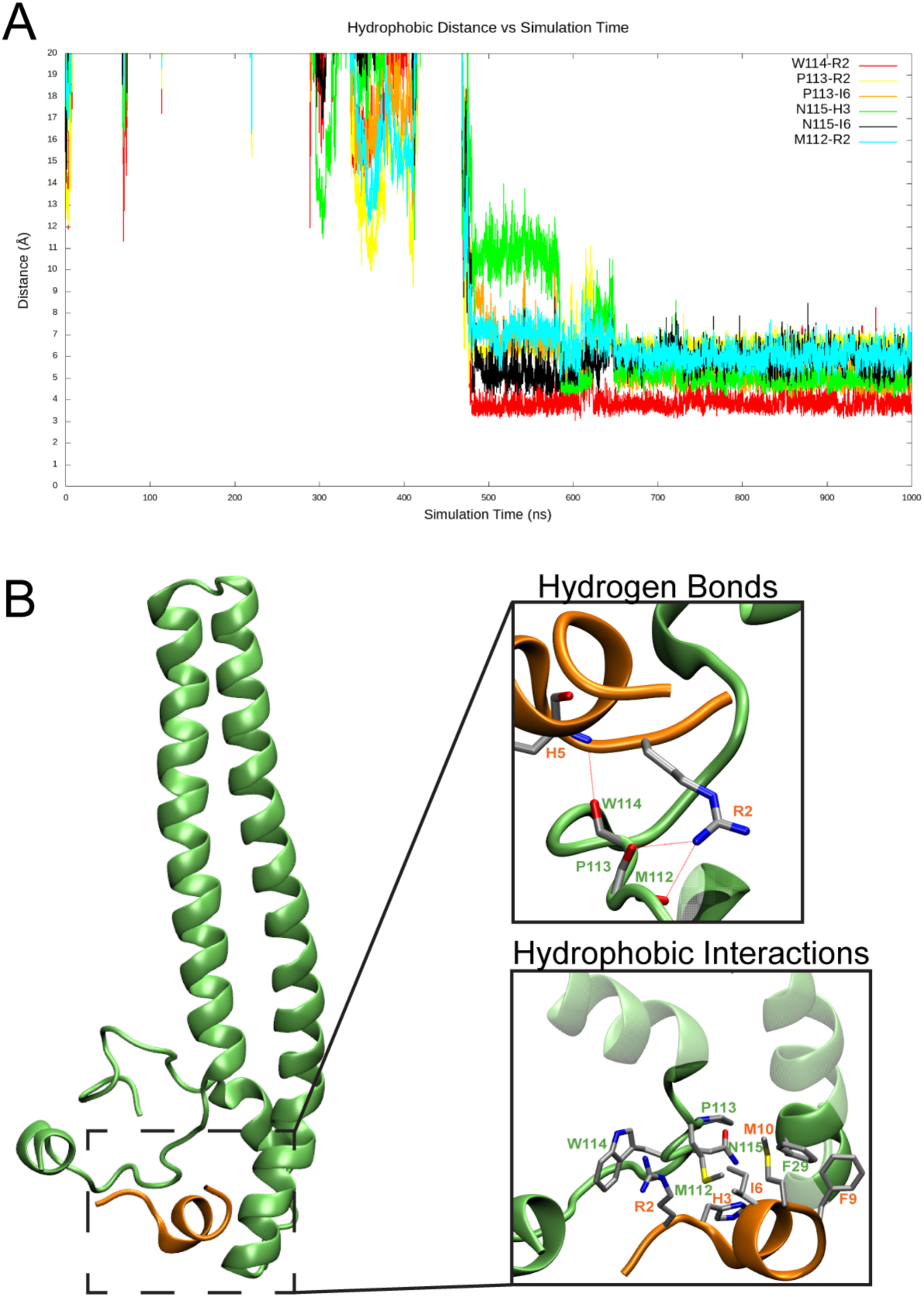
Braftide interacts with CDC37 independently. A) Distance plot of braftide with CDC37 along the MD trajectory. B) CDC37 interaction with braftide and zoomed in view of specific residues interacting via hydrogen bonding (above) and through hydrophobic stacking (below).

**Figure S8.**
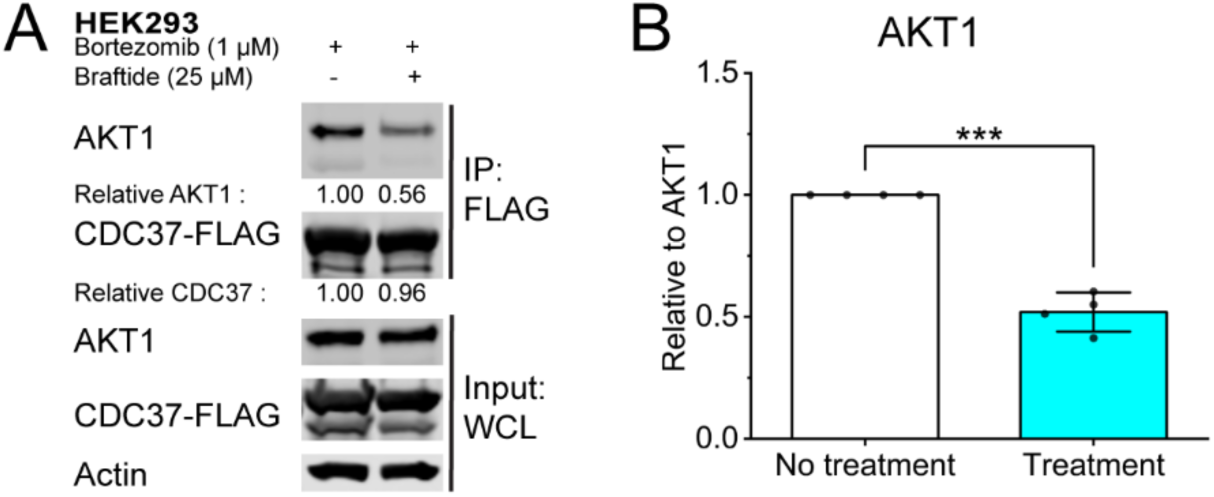
Braftide treatment disrupts client AKT1 from CDC37-FLAG in HEK293 cells. A) Representative immunoblot of immunoprecipitated CDC37 with co-immunoprecipitated AKT1 of three biological replicates in the absence and presence of Braftide (25 µM) treatment. Cells are bortezomib pretreated to prevent proteasomal degradation. B) Densitometry analysis of four biological replicates (n=4). Graph bars represent the mean ± SD with corresponding P-values (*P<0.05, **P<0.01, ***P<0.001). Statistical significance was determined via one-way ANOVA, followed by the post-hoc Tukey’s test.

**Figure S9.**
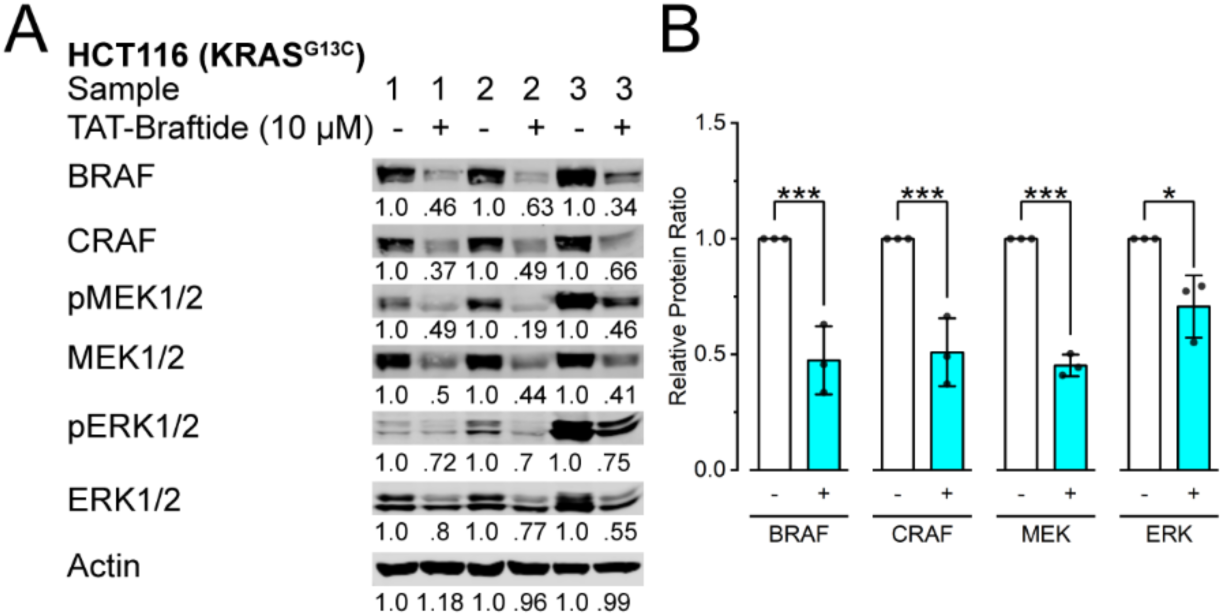
Braftide decreases MAPK pathway protein levels in samples prepared for global proteomics. A-B) Immunoblot of TAT-Braftide-mediated downregulation of RAF-MEK1/2-ERK1/2 pathway protein levels in HCT116 cells (constitutively active KRASG13C mutant cancer cell line) (n=3). KRAS G13C hyperactivates the RAF-MEK1/2-ERK1/2 pathway. These three biological replicates were subjected to LC-MS/MS for differential protein expression in the presence and absence of Braftide treatment (10 µM, 1 hr, 37°C). The relative ratio is relative to the non-treatment sample (no treatment/Braftide treatment). B/CRAF, phosphorylated MEK1/2 (pMEK1/2), total MEK1/2, phosphorylated ERK1/2 (pERK/2), total ERK1/2, and actin are visualized in the immunoblots. B) Densitometry analysis of the samples used for differential global proteomic identification of the MAPK pathway constituents. Statistical significance was determined via two-way ANOVA, followed by the post-hoc Tukey’s HSD (honest significant difference) test. Graph bars represent the mean ± SD with individual data points per biological replicate with corresponding P-values (*P<0.05, **P<0.01, ***P<0.001).

**Figure S10.**
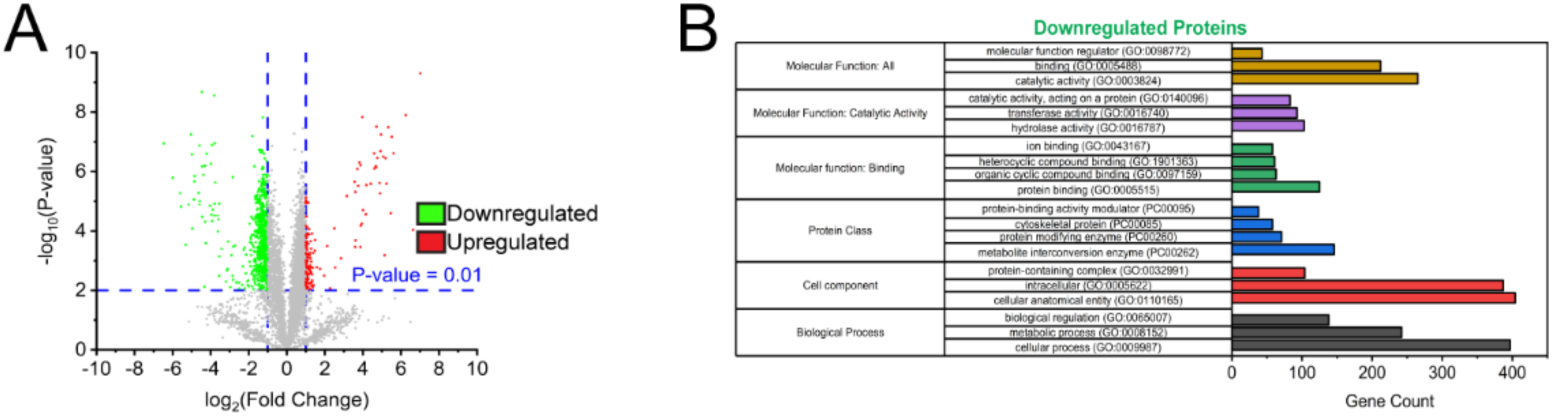
Braftide downregulates endogenous kinase levels. A) Volcano plot of 5405 proteins as a result of Braftide treatment identified in three biological replicates. The blue vertical lines represent the 2-FoldChange (FC) cutoff. Neon green dots represent the 627 proteins that are downregulated. Red dots represent the 118 upregulated proteins. B) Gene ontology (GO) analysis of downregulated proteins in which the top 3-4 most Braftide treatment enriched categories are graphed by gene count (frequency) using the Panther GO database.

**Figure S11.**
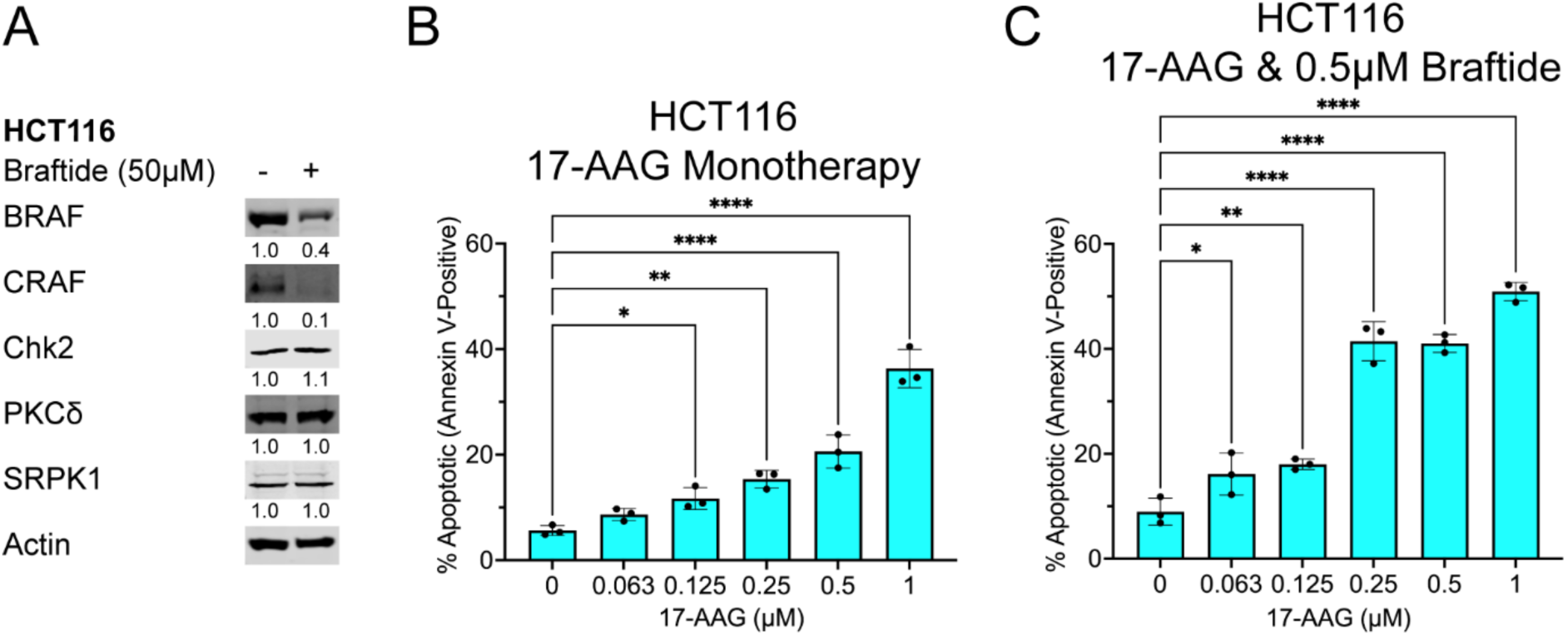
**Braftide specifically affects kinases reliant on HSP90-CDC37**. A) Representative immunoblot of selected kinases that are unaffected by Braftide treatment. B-C) Braftide synergizes with 17-AAG in HCT116 cells to trigger cell death at lower treatment concentrations. Cells were stained with Annexin V, an apoptosis marker, and sorted via flow cytometry (n=3).

Supplementary Information. Mass spectrometry data from Braftide treated HCT116 cells with differential protein expression (xls file)

